# Scalable Solar-Driven Chemical Production by Semiconductor Biohybrids Synthesized from Wastewater Pollutants

**DOI:** 10.1101/2023.02.28.530441

**Authors:** Shanshan Pi, Wenjun Yang, Wei Feng, Ruijie Yang, Weixiang Chao, Wenbo Cheng, Lei Cui, Zhida Li, Yiliang Lin, Chen Yang, Lu Lu, Xiang Gao

## Abstract

Semiconductors biohybrids integrate the best of biological catalysts and semiconductor nanomaterials for solar-to-chemical conversion. To realize the potential of hybrid systems at the commercial level, it remains an urgent need for cost-competitive and environmentally friendly approaches to scaling up. Here, we successfully tackle this challenge through developing biohybrid route that co-utilize multi-pollutants in wastewater to produce semiconductor biohybrids *in-situ* for solar-to-chemical production. To achieve cost-effective biohybrid production, we introduced an aerobic sulfate reduction pathway into *Vibrio natriegens* to enable the direct utilization of the heavy metal ions (*i*.*e*., Cd^2+^), sulfate, and organics in the wastewater to biosynthesize functional semiconductor nanoparticles in living *V. natriegens*. Furthermore, 2,3-butanediol biosynthetic pathway was introduced into the *V. natriegens* hybrid to couple the solar energy for enhanced bioproduction. We demonstrated the scalability of this system in a 5-L illuminated fermenter using wastewater as the feedstock, which resulted in production of 13 g/L of 2,3-butanediol. Life cycle analysis showed this specific biohybrid route had a significantly lower cost and reduced CO_2_ emission compared to both pure sugars fermentation and fossil-based routes. In addition to providing a promising step toward sustainable commercializing semiconductor biohybrids for biomanufacturing, our work may lead to hybrid living matter toward future waste to wealth conversion.

## Introduction

Biomanufacturing with sugar fermentation offers an opportunity to switch from fossil fuels to renewable energy sources, which contributes to carbon emissions reduction and climate changes alleviation (Fig. 1a)^1,2^. In the state-of-the-art biomanufacturing, the sugar substrates are oxidized in the process to generate reducing-energy (NAD(P)H), which releases CO_2_ and thus reduces carbon yield during chemical production^1,3^. Recently, the semiconductor biohybrids offer a promising strategy in biomanufacturing through decoupling carbon and bioproduction of reducing-energy^4,5^. Through integrating the efficient light-harvesting materials with microbial cell factories, the biohybrids can capture light energy and produce reducing-energy, reducing/eliminating the loss of carbon (Figs. 1a and 1b)^4,5^. Photosynthetic semiconductor biohybrids integrate the best attributes of biological whole-cell catalysts and semiconducting nanomaterials, and it serves a novel approach to utilize solar energy for chemical production^6^. Currently, there are limited applications of semiconductor hybrids due to the relatively high cost of large-scale biohybrids constructions^7,8^. On one hand, traditional semiconductors synthesis through physical or chemical methods is uneconomic and environmentally unsustainable^9^. On the other, biosynthesis of semiconductor nanoparticles, such as cadmium sulfide, via microorganism typically involves highly expensive cysteine precursor^4,10,11^. It remains a major challenge to leverage biohybrids to achieve scalable chemical production in a cost-effective and environmentally sustainable manner.

**Fig. 1.**
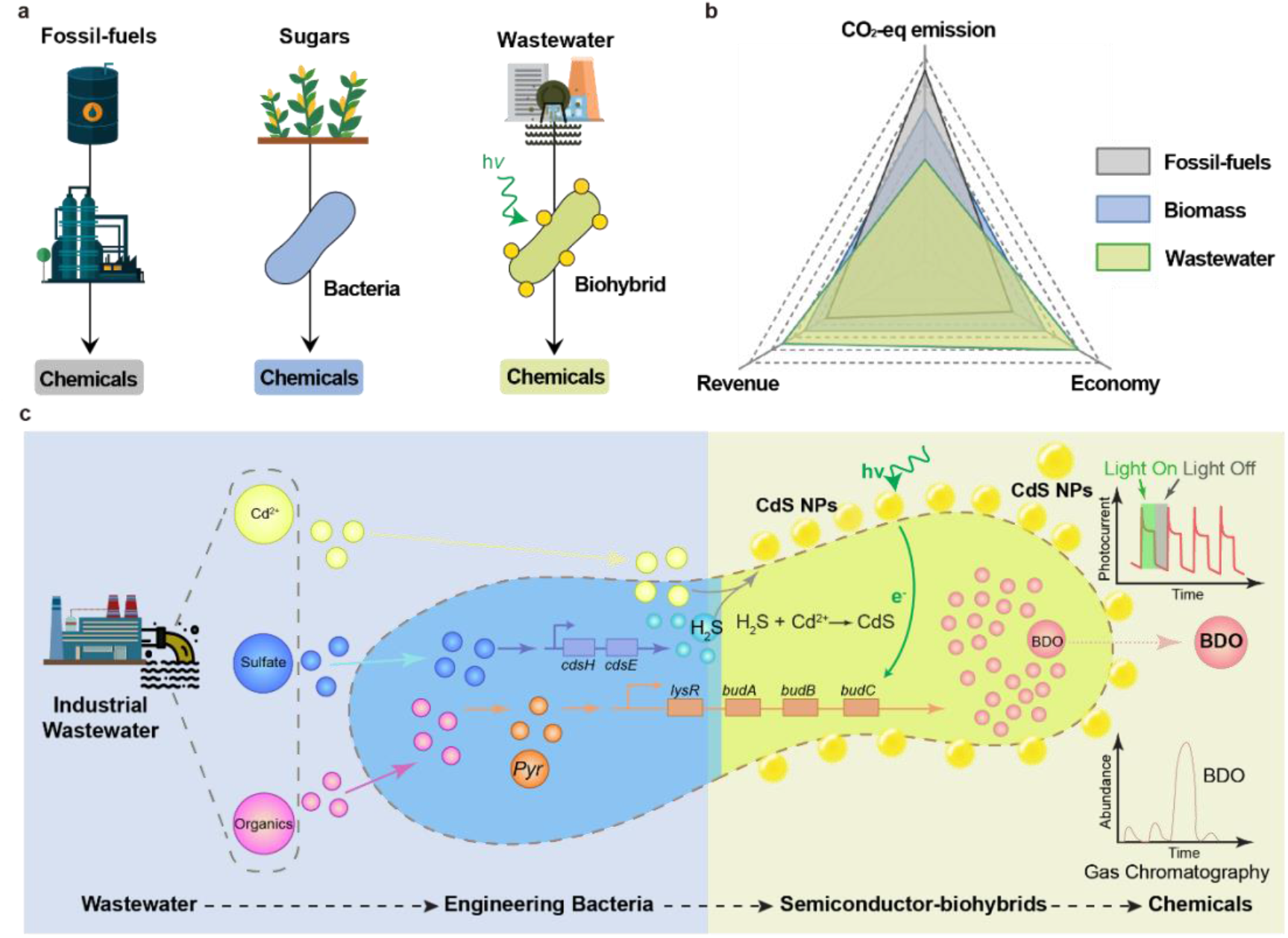
Schematic diagram of sustainable production of semiconductor biohybrids for *in-situ* solar-driven chemical production from multi-pollutants in wastewater using engineering *V. natriegens*. **a**) Schematic of chemicals production by refinery of fossil-fuels, pure bacterial fermentation of sugars, and biohybrids synthesis using wastewater. **b**) Comprehensive evaluation of sustainability of these routes by CO_2_ equivalent of greenhouse gas (GHG) emissions, economy of cost advantages, and revenue from extended by-products and environmental remediation. **c)** The effluent of industrial wastewater contains pollutants including Cd^2+^, sulfate and organics. These pollutants could be co-utilized by engineering *V. natriegens* to construct biohybrid system for solar-driven chemical production *in-situ*.

The waste-to-wealth approach is well aligned with renewable energy and eco-friendly sustainability^12,13^. In recent years, sustainable biomanufacturing has demonstrated the possibility of making a range of commodity chemicals from waste, including plastic^14^, food waste^15^ and industrial gas^16^. Recently, an engineered sulfide-producing yeast was constructed to precipitate heavy metals in sulfate-amended wastewater by forming metal sulfide as potential semiconductor for environmental remediation^17^. Thus, it is a promising approach to directly use sulfate and heavy metal containing wastewater as feedstock towards biohybrids synthesis for solar-to-chemical production, turning the waste into wealth. Additional, industrial wastewaters typically contain organic wastes (e.g. carbohydrate wastes), which are generally environmental pollutants and require costly remediation^18,19^, but could be utilized for bacterial growth and biochemical production^2,18^. Hence, we propose that an engineered microorganism that could efficiently co-utilize sulfate, heavy metals, and carbon resources in wastewater simultaneously for biohybrids synthesis, if successfully developed, will pave a new way to advanced waste-to-wealth environmental sustainability as well as biomanufacturing with higher carbon yield.

Here, we design and engineer *Vibrio natriegens* to synthesize semiconductor biohybrids directly from real industrial wastewater and further produce a commodity chemical in 5 liters scale (Fig. 1c). *Vibrio natriegens*, marine bacterium, is a promising industrial production host with fast growth rate (doubling time < 10 min), high substrate consumption rates^20^ and exceptional tolerance to high salt concentration which is a common occurrence in wastewater^21,22^. Besides, *V. natriegens* could utilize more than 60 different carbon sources, including most the carbohydrates, proteins, carboxylic/ aromatic acid and alcohols, which are the common organics in the wastewater^19,23^. Thus, the *V. natriegens* could be a great candidate to utilize wastewater for solar-to-chemical production through turning waste into wealth. By introducing an aerobic sulfate reduction pathway into *V. natriegens*, the engineered strain could produce hydrogen sulfide (H_2_S) directly from sulfate-containing wastewater, and further biomineralize the metal ions to form biohybrids *in situ*. We find this engineering could directly utilize real industry wastewater (organic wastewater combined with electroplating wastewater) for constructing metal sulfide biohybrids, confirmed by electron microscopy imaging, X-ray diffraction (XRD) and inductively coupled plasma mass spectrometry (ICP-MS). We also confirm the wastewater-derived biohybrid have a good efficiency for light absorption and photoelectrons generation through photoelectrochemical analysis. We further design and introduce a 2,3-butanediol (BDO) biosynthetic pathway to enable the biohybrid to produce BDO from organics in wastewater. The resulting biohybrid cells increase intracellular reduced nicotinamide adenine dinucleotide (NADH) under light condition and facilitate higher BDO production and higher conversion yield compared with pure bacterial cells (Fig. 1c).

In lab-scale fed-batch fermentation, 13.09 g/L of BDO is achieved by engineered biohybrids constructed from real industrial wastewater, which enables *in-situ* scalable solar-driven chemical production with waste organics serving as the carbon source and sacrificial agents of photogenerated hole. Life cycle analysis (LCA) confirm a lower greenhouse gas (GHG) emissions and lower cost for product with our developed platform compared to both pure bacterial fermentation and fossil-based routes (Fig. 1b). In this study, we develop a cost-competitive and environmentally friendly approach that is promising for sustainable production of semiconductor biohybrids towards solar-driven chemical production utilizing multi-pollutants in wastewater.

## Results

### Engineering *V. natriegens* to synthesize semiconductor biohybrid

The organic carbon, heavy metal ions, and sulfate in wastewater can serve as major substrates for semiconductor biohybrids construction and solar-to-chemicals production (Fig. 1c). In order to achieve biohybrids construction in wastewater and waste-to-wealth sustainability, our first step was to engineer *V. natriegens* so that it produced sulfide directly from sulfate, a common pollutant in wastewater^24^, rather than adding expensive cysteine precursors. The conversion of sulfate to H_2_S is found in nature by a diverse group of sulfate-reducing bacteria (SRB), but they have a slow growth rate and typically require strict cultivation conditions because they are obligate anaerobes^25^. Moreover, those SRB usually metabolize simple organics to reduce sulfate^25,26^. Alternatively, it is possible to introduce aerobic sulfide reduction pathway into *Escherichia coli* to utilize sulfate to overproduce cysteine, which was then converted to H_2_S by cysteine desulfhydrase^27^. Inspired by this, we introduced a pathway contains a mutant serine acetyltransferase (CysE) that is insensitive to feedback inhibition by cysteine, and cysteine desulfhydrase (CdsH) into *V. natriegens* (Fig. 2a), resulting in an aerobic strain of XG203. To examine the capability to produce H_2_S, we cultured the XG203 in a well-defined minimum medium with sulfate supply. XG203 produced 106 ppm H_2_S in 24 hours and the average production rate reached about 4.42 ppm/h (Figs. 2b). Meanwhile, there was no detectable H_2_S produced even after 24 hours by wild-type *V. natriegens* (Fig. 2c), confirming the successful introduction of aerobic sulfide production pathway.

**Fig. 2.**
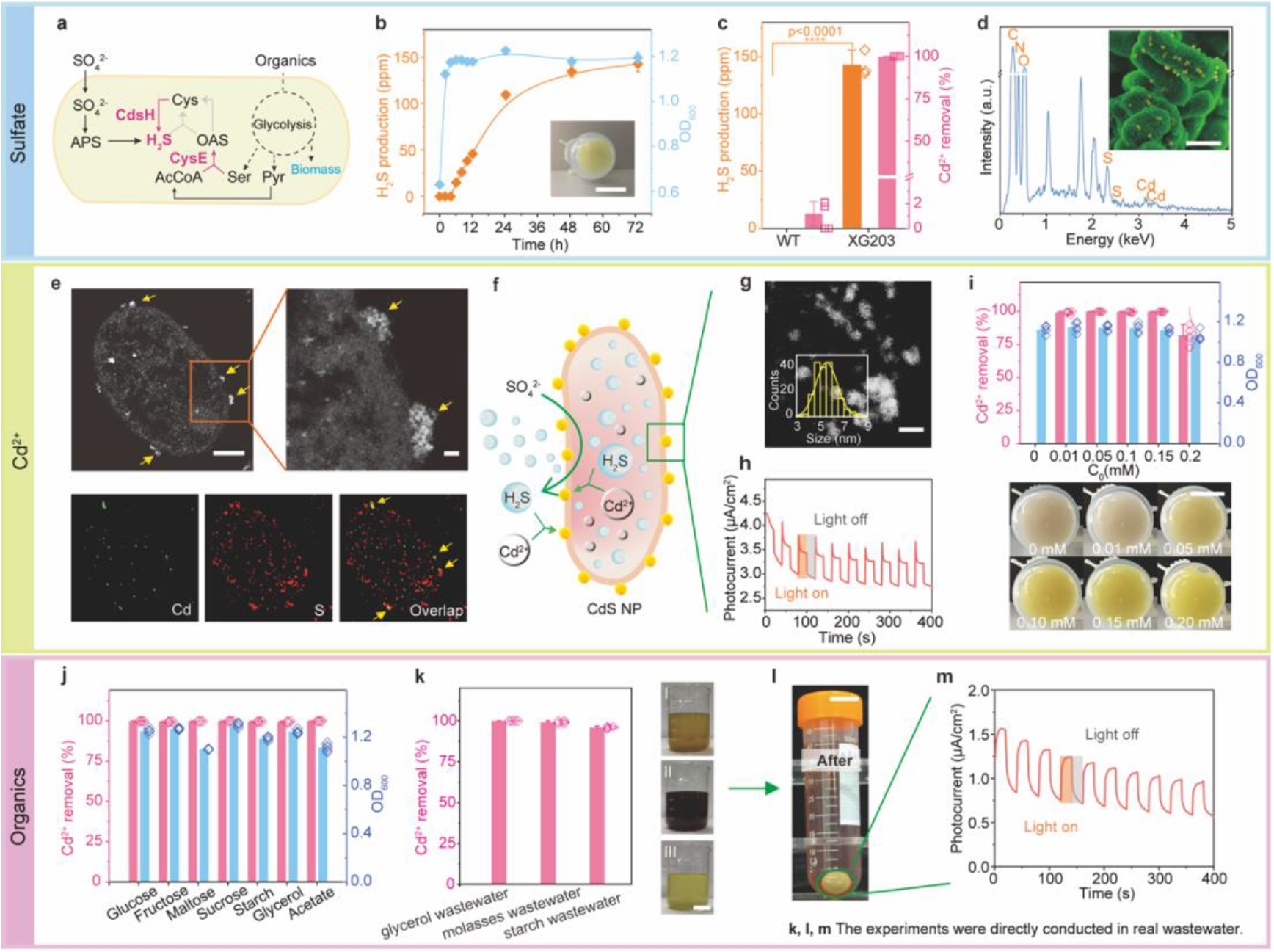
Design *V. natriegens* to produce semiconductor biohybrid from wastewater. **a**) Engineering an aerobic SO_4_ ^2-^ assimilation pathway in *V. natriegens* to produce H S. The CysE increases and CdsH were overexpressed. OAS, O-acetylserine; Pyr, pyruvate; Ser, serine; AcCoA, Acetyl-CoA; APS, Adenosine 5’-phosphosulfate; Cys, cystein. **b**) The H_2_S production and cell growth of strain XG203 in a well-defined minimum medium. Error bar: standard deviation, n≥3. The inset is the pellets of strain XG203 after culturing with Cd^2+^. Scale bar, 2 cm. **c**) Comparison of Cd^2+^ removal and H_2_S production between wild-type and engineering *V. natriegens*. Error bar: standard deviation, n≥3. **d**) SEM image (inset) and EDS mapping of biohybrid sample collecting from **c**. Scale bar, 0.5 μm. **e**) Cross-sectional STEM image of biohybrid sample collecting from **c**. Scale bars, 0.2 μm (left) and 20 nm (right). **f**) Schematic diagram of *in-situ* production of semiconductor biohybrid. **g**) HAADF-STEM image of isolated CdS nanoparticles. High-resolution transmission electron microscopy (HRTEM) images are used to analyze the size of nanoparticles (more than 200 particles). Scale bar, 5 nm. **h**) A representative photocurrent curve of isolated nanoparticles with light on/off cycles (20 s on and off, 100 mW/cm^2^). The isolated nanoparticles were lysed from biohybrids of **c. i**) The stain XG203 was cultured in a well-defined minimum medium with sulfate and different concentration of Cd^2+^ (0.01∼0.2 mM) for biohybrid construction. Error bar: standard deviation, n=5. The lower images are pellets of strain XG203 after culturing under different Cd^2+^ concentrations. Scale bar, 1.5 cm. **j**) The removal of Cd^2+^ and cell growth by strain XG203 after culturing in a well-defined minimum medium with those common carbon sources from wastewater. Error bar: standard deviation, n≥4. **k**) The removal of Cd^2+^ by strain XG203 after culturing three kinds of real wastewater. Error bar: standard deviation, n=4. Three wastewaters contain glycerol wastewater (I), molasses wastewater (II), and starch wastewater (III) as carbon sources, respectively. Visual representation of corresponding wastewaters (right). Scale bar, 5 cm. **l**) The pellets of engineering strain after culturing in wastewater with organics supply by molasses wastewater collecting from **k**. Scale bar, 1.5 cm. **m**) A representative photocurrent curve of biohybrid samples collecting from **l**. P values are determined by a two-tailed unpaired t-test.

We further cultured the strain XG203 in a well-defined minimum medium with sulfate and 0.1 mM CdCl_2_ to investigate its capability to utilize the heavy metal resources with the production of H_2_S. Impressively, almost all the Cd^2+^ (99.91%) were removed within ∼ 2 hours with a removing rate of 4.56 mg/L/h (Fig. S1a). The bacterial culture became yellow simultaneously, suggesting the formation of CdS nanoparticles (Fig. 2b). Meanwhile, the wide-type *V. natriegens* remained white color and only removed 1.19% Cd^2+^ from medium even after 24 hours (Fig. 2c). The small portion of Cd^2+^ removal is most likely due to simple adsorption^28^.

In order to confirm the yellow pellets were CdS-bacteria hybrids, we first visualized them with electron microscopy. Scanning electron microscope (SEM) showed that bacterial cells in the hybrids deposit nanoparticles with an average size of ∼55.11 nm *in-situ* (Fig. 2d and Fig. S1b), while there is no obvious nanoparticles observed in *V. natriegens* cultured in medium without Cd^2+^ (Fig. S1c). The energy dispersive spectroscopy (EDS) mapping confirmed that those particles are composed of Cd and S elements (Fig. 2d). Furthermore, we prepared the cross-sectional samples through microtoming. Scanning transmission electron microscopy (STEM) further confirmed those particles mainly locate on the surface of bacterial cells (Fig. 2e), and STEM energy dispersive spectroscopy (STEM-EDS) mapping showed that the particles are also composed of Cd and S elements (Figs. 2e and 2f). We speculate the nanoparticles that deposit on bacterial cells are CdS NPs.

Afterwards, we lysed the biohybrids and collected the nanoparticles for further characterization. The nanoparticles have a good crystallinity with the main lattice spacing of 3.36 Å and 2.05 Å (Figs. S1d and S1e), and the X-ray powder diffraction (XRD) patterns confirmed those nanoparticles are CdS (Fig. S1f). High-resolution X-ray photoelectron spectroscopy (XPS) of Cd 3d also confirmed the chemical state of the cadmium species is Cd (II) in the nanoparticles (Fig. S1g). STEM analysis revealed these dispersive nanoparticles have an average size of ∼5.54 nm which is similar to previous report of extracted nanoparticles in microorganism^9,29^ (Fig. 2g). Notably, the size of lysed particles is much smaller than the particles deposited on the cell surfaces, indicating aggregation of multiple particles (∼ 10) during biosynthesis. High-angle annular dark-field STEM (HAADF-STEM) and selected area electron diffraction (SAED) of the particles indicated a polycrystalline structure (Fig. 2g and Fig. S1d). To discern the function of the biomineralized CdS NPs, we performed UV-Vis spectrum and electrochemical characterization to measure photoelectrochemical performance of particles. The CdS nanoparticles have a direct band energy of about 2.58 eV (Fig. S1h), and the photo-induced current was about 0.53 μA/cm^2^ (Fig. 2h), which is close to that reported range in previous reports^11,30^.

The concentrations of heavy metal ions, such as Cd^2+^, in common industrial wastewater have a wide range from several ppm to the thousands of ppm^31-33^. For high concentration wastewater, a pretreatment based chemical/physical (e.g. precipitation and adsorption) approaches is generally set within factories by mandatory regulations to remove most of metals for reducing the environmental risks^34^, which leads to an effluent with relative low metal ion concentration (below dozens of ppm) that is compatible to sustainable biological treatment^31,35^, such as that in domestic wastewater treatment plant. To confirm the capability of strain XG203 to form biohybrids from the majority of wastewater resources, we cultured strain XG203 with Cd^2+^ from 0.01 mM to 0.2 mM (1-22 ppm) as a stress testing. Surprisingly, the engineering strain removed almost all the Cd^2+^ (above 99%) when the concentration no more than 0.15 mM, and removed 81.92% of Cd^2+^ with negligible inhabitation of bacterial growth even in a very high concentration (0.2 mM Cd^2+^) (Fig. 2i). Also, with the increase of Cd^2+^ concentration in the medium, the culture become more yellow, which is consistent with the increase of biohybrids and CdS nanoparticles formation (Fig. 2i). This strategy can be extended to other heavy metals as well, and heavy metal-contaminated wastewaters are feedstocks for hybrid construction with benefits of effective remediation (Fig. S2 and Table S1). The universality of our developed strategy was further demonstrated by other metallic sulfides, such as PbS (Fig. S3) and HgS (Fig. S4).

### Engineering *V. natriegens* to utilize different carbon source in wastewater

Aside from the different species and concentrations of heavy metal ions in industrial wastewater, organics can also vary significantly^19,23^. To establish a sustainable and economical chemical production process, it requires the strain XG203 to be able to adapt to different organics in wastewater resources. The major organic component in different industrial wastewater varies (detail description in Supplementary Method). For instance, wastewater generated during sugar production mainly include molasses, while those generated for biodiesel production generally contain glycerol^36,37^. Molasses waste is a complex of sucrose, fructose and glucose according to the high-performance liquid chromatography (HPLC) analysis (Table S2). To test the broad spectrum of substrates utilization, we cultured strain XG203 in a well-defined minimum medium (amendment by sulfate and Cd^2+^) with different organics, including glucose, fructose, maltose, sucrose, starch, glycerol, and acetate, and the strain XG203 can all grow very well with high Cd^2+^ (all above 99%) removing efficiency and biohybrids formation (Fig. 2j and Fig. S1h).

We further utilized the real industrial wastewater (detailed compositions in Table S4), which was prepared by mixing organic wastewater (generation from biodiesel, corn starch or sugar industries with detailed compositions in Table S2) with electroplating wastewater (detail compositions in Table S3). The strain could all synthesize biohybrids and simultaneously achieve high Cd^2+^ removing efficiency of 95.93∼99.92% (Fig. 2k). The bacterial culture turned yellow over time, indicating the generation of CdS biohybrids using wastewater (Fig. 2l). We further collected the biohybrids directly synthesized from real industrial wastewater with organics supply by molasses wastewater for photoelectrochemical measurement, and the biohybrids show the photo-induced current of ∼0.51 μA/cm^2^ (Fig. 2m), which is close to the photocurrent of previous reported biohybrids^38^.

Taking together, with our tailored and engineered strain, we are able to synthesize semiconductor biohybrids directly from wastewater with successful utilization of the heavy metals, sulfates, and carbon in wastewater, paving a promising path for large-scale production for biomanufacturers.

### Solar-to-chemical production by the semiconductor biohybrids from wastewater

After confirming the successful synthesis of biohybrids using real wastewater resources, we further tested whether these biohybrids could perform solar-to-chemical production *in situ* for renewable energy and sustainable biomanufacturing. 2,3-butanediol (BDO) is a promising bulk chemical for a wide range of potential utilization^39^. Therefore, we introduced the BOD biosynthetic pathway into *V. natriegens* to demonstrate the capability of our biohybrids for solar-to-chemical production. The BDO biosynthetic pathway contains three enzymes (Fig. 3a). Acetolactate synthase (ALS) and acetolactate decarboxylase (ALDC) convert pyruvate to acetoin which is catalyzed by 2,3-butanediol dehydrogenase (BDH) and converted to BDO by using the reducing-power NADH^40,41^ (Fig. 3a). We cloned the BDO pathway gene cluster from *Enterobacter cloacae* and expressed in strain XG203 under the control of a native promoter^42^, and the resulting strain was designed as XG203A.

**Fig. 3.**
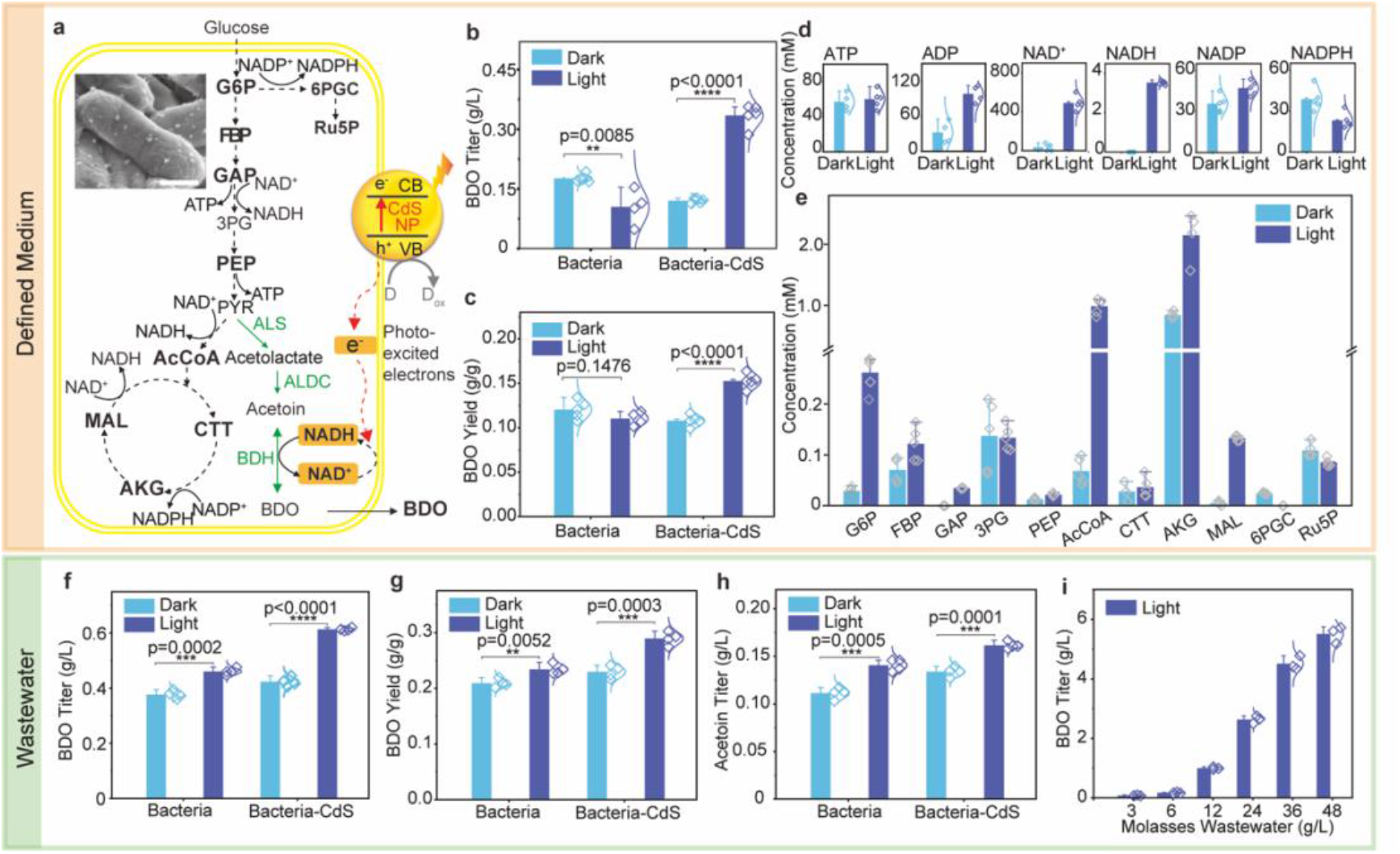
Solar-to-chemical production by the semiconductor biohybrids from wastewater. **a**) A solar-enhanced BOD biosynthetic pathway in semiconductor biohybrid. G6P, Glucose 6-phosphate; FBP, Fructose 1,6-bisphosphate; GAP, Glyceraldehyde 3-phosphate; 3PG, 3-phosphoglycerate; PEP, Phosphoenolpyruvate; PYR, pyruvate; AcCoA, Acetyl-CoA; CTT, Citrate; AKG, α-ketoglutarate; MAL, Malate; 6PGC, 6-phosphogluconate; Ru5P, Ribulose 5-phosphate; CdS NP, cadmium sulfide nanoparticles; CB, conduction band; VB, valence band; D, electron donor; D_ox_, oxidized donor. SEM image (inset) of semiconductor bihybrid. Scale bar, 0.5 μm. **b**) BDO production by pure bacterial system only and bacteria-CdS hybrid system under dark and light condition (4.2 mW/cm^2^). Error bar: standard deviation, n≥4. **c**) Glucose-to-BDO conversion yield of bacteria and bacteria-CdS biohybrid. Error bar: standard deviation, n≥4. **d**) Concentration of intracellular cofactors in semiconductor biohybrid under dark and light condition (4.2 mW/cm^2^). Error bar: standard deviation, n≥4. **e**) Concentration of intracellular metabolites of central carbon metabolism in semiconductor biohybrid under dark and light condition (4.2 mW/cm^2^). Error bar: standard deviation, n≥3. **f**) BDO production of pure bacterial system and biohybrid system using wastewater under dark and visible light condition (4.2 mW/cm^2^). Error bar: standard deviation, n=4. **g**) Waste sugar-to-BDO conversion yield of bacteria and biohybrid system using wastewater. Error bar: standard deviation, n=4. **h**) Acetoin production of bacteria and biohybrid system using wastewater under dark and visible light condition (4.2 mW/cm^2^). Error bar: standard deviation, n=4. **i**) Solar-driven BDO production by semiconductor biohybrids using wastewater containing different concentration of molasses (3∼48 g/L). Error bar: standard deviation, n=3. P values are determined by a two-tailed unpaired t-test.

To test the successful integration of BDO pathway and explore the biological mechanism, we firstly tested the ability of strain XG203A for light-driven chemical production in a well-defined minimum medium (modified M9 medium with 5 g/L glucose, Cd^2+^ and sulfate). We confirmed illuminated biohybrid system can produce BDO with a titer of 0.33 g/L, showing 1.90 times improvement compared to bacterial system (Fig. 3b). Also, the glucose-to-BDO conversion yield increase from 0.12 g/g to 0.15 g/g, with an increasement of 26.69% (Fig. 3c), likely due to the regeneration of additional NADH from illuminated CdS nanoparticles (see more discussion below). We also found the BDO production and yield of biohybrid system was 2.78 and 1.41 times than that of the dark condition (Fig. 3b and 3c). However, the strain XG203A without CdS nanoparticles, the BDO production and yield under light was 40.76% and 8.39% lower than the dark condition. The experiment suggested that biohybrid system could be used for efficient biomanufacturing as previous reports^43-45^.

To explore the possible factors contributing for increasing solar-to-chemical production, we collected CdS-biohybrid samples from a well-defined minimum medium under light and dark condition, and performed targeted metabolites quantification by using liquid chromatography-tandem mass spectrometry (LC-MS/MS). We firstly analyzed the effect of cofactor regeneration induced by photo-induced electron^5,8^ and found NAD^+^ and NADH were increased to 490.58 μM and 3.54 μM in hybrids under light condition, which is much higher than that of dark condition (Fig. 3d). While, NADPH concentration in light condition is 40.41% lower than dark and ATP concentration is similar in light and dark condition (Fig. 3d). In bacterial cells, the central carbon metabolism often couples with the energy metabolism^6,46^, we further analyzed the regulation of photo-induced electrons on central carbon metabolism. NADH is mainly produced from the glycolysis pathway (Embden-Meyerhof-Parnas, EMP pathway) and tricarboxylic acid cycle (TCA), while NADPH is synthesized from pentose phosphate pathway (PPP) (Fig. 3a). We found glucose 6-phosphate (G6P) and fructose 1,6-bisphosphate (FBP) in MEP pathway, and citrate (CTT) and a-ketoglutarate (AKG) in TCA cycle were increased in light condition compared with its counterpart with dark (Fig. 3e). Correspondingly, the 6-phosphogluconate (6GPC) and ribulose 5-phosphate (Ru5P) in PPP were lower in light condition than that of dark (Fig. 3e). Those results suggest that the central carbon metabolism also contributes to the NADH increase in the biohybrid system. In addition, we found acetoin, a direct precursor for BDO production, increased by 84.33% in light condition (Fig. S5a). The higher concentration of substrates, acetoin and NADH, could provide a driven force for BDO production.

Taken together, those results suggest illuminated semiconductor biohybrid system could stimulate the activity of intracellular metabolism to provide more reducing power and substrate for final chemical production, which was also found in previous research^5,46^.

After confirming the successful production of BDO with strain XG203A and probing the mechanism behind the photosynthesis, we further applied the strain XG203A in real wastewater containing 4.1 g/L sugar, Cd^2+^ and sulfate. As shown in Fig. 3f, the BDO production of illuminated biohybrid system using wastewater was superior to all other conditions, with a titer of 0.61 g/L, showing 1.45-fold increase compared with its counterpart under dark and 1.33 times increase compared with engineering bacteria in absence of nanoparticles under light. Waste organics (sugar)-to-BDO conversion yield was also increased 26.10% compared with dark condition of biohybrid system (Fig. 3g). Acetoin production using wastewater by illuminated biohybrid system was also the highest compared to all other conditions (Fig. 3h and Fig. S5b). Moreover, the production of BDO was gradually increased by adding higher concentration of molasses wastewater in the illuminated biohybrid culture, reaching to 5.51 g/L with 48 g/L molasses (Fig. 3i). The experiment suggested that wastewater-based biohybrid system could be *in-situ* used for efficient solar-to-chemical production, with an increased carbon conversion into chemicals.

### Scale-up chemical production with biohybrids in wastewater

Finally, to demonstrate the performance of biohybrids in the scalable production by co-utilization of multi-pollutants in wastewaters, we tested solar-driven BDO production by the *in-situ* production of semiconductor biohybrids using our engineered strain XG203A with a pH-controlled fed-batch 5-L fermenter under illumination (Fig. 4a). XG203A utilized 78.05% of sugars from the wastewater for *in-situ* production of semiconductor biohybrids and chemicals (Fig. 4b). The production of BDO and acetoin reached 13.09 g/L and 1.13 g/L, respectively (Fig. 4c), and the maximum productivity of BDO was 0.67 g/L/h (Fig. 4d).

**Fig. 4.**
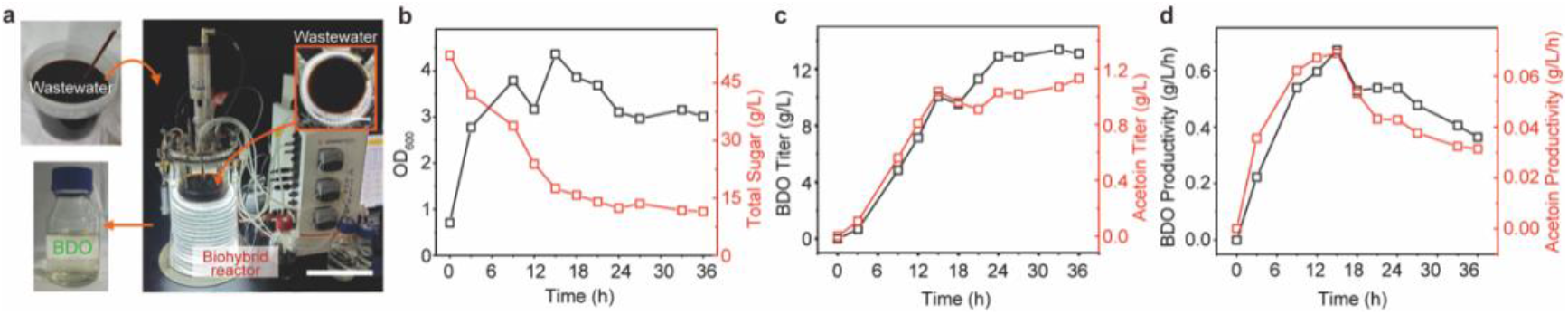
Scale-up chemical production of biohybrid system using real wastewater. **a**) Visual representation of the 5-L illuminated bioreactor for BDO production by *in-situ* production of semiconductor biohybrid using wastewater. Wastewater was converted to valuable chemical (BDO) by strain XG203A in the illuminated bioreactor. Scale bars, 20 cm and 6.5 cm (inset). **b**) Time courses of sugar consumption and growth (OD_600_) of semiconductor biohybrid system during BDO production. **c**) Titer of BDO and acetoin produced by semiconductor biohybrid. **d**) Productivity of BDO and acetoin produced by semiconductor biohybrid.

We used life cycle analysis (LCA) to compare the BDO production by lab-scale wastewater-based biohybrids with current production from fossil-fuels and bacterial fermentation of sugars (Fig. 5a). In order to quantify the practical effects of biohybrid synthesis, a model used for evaluation of industrial fermentation is modified and employed here to overcome the uncertainties in the lab-scale level (Fig. 5b). Both the GHG emission and cost of BDO by biohybrid (0.50±0.02 kgCO_2_-eq/kg BDO, 0.39±0.04 $/kg BDO) at the 95% confidence level are superior than those of fossil-fuels production (1.25±0.002 kgCO_2_-eq/kg BDO, 0.81±0.03 $/kg BDO) and pure sugar fermentation (0.82±0.003 kgCO_2_-eq/kg BDO, 0.48±0.02 $/kg BDO) (Figs. 5c and 5d). The lower GHG emission and cost are attributed to utilize wastewater as feedstock for reducing the requirement for wastewater treatment and recover CdS nanoparticles as revenues for integrating solar energy during biomanufacturing (Table S12). It should be noting several factors which not considered in our model, such as the sacrificial agents, the different separation efficiency of photo-induced electron-hole pairs, and the competition between complex organics and metal ions. Further work is need for a more accurate LCA of biohybrid routes using wastewater.

**Fig. 5.**
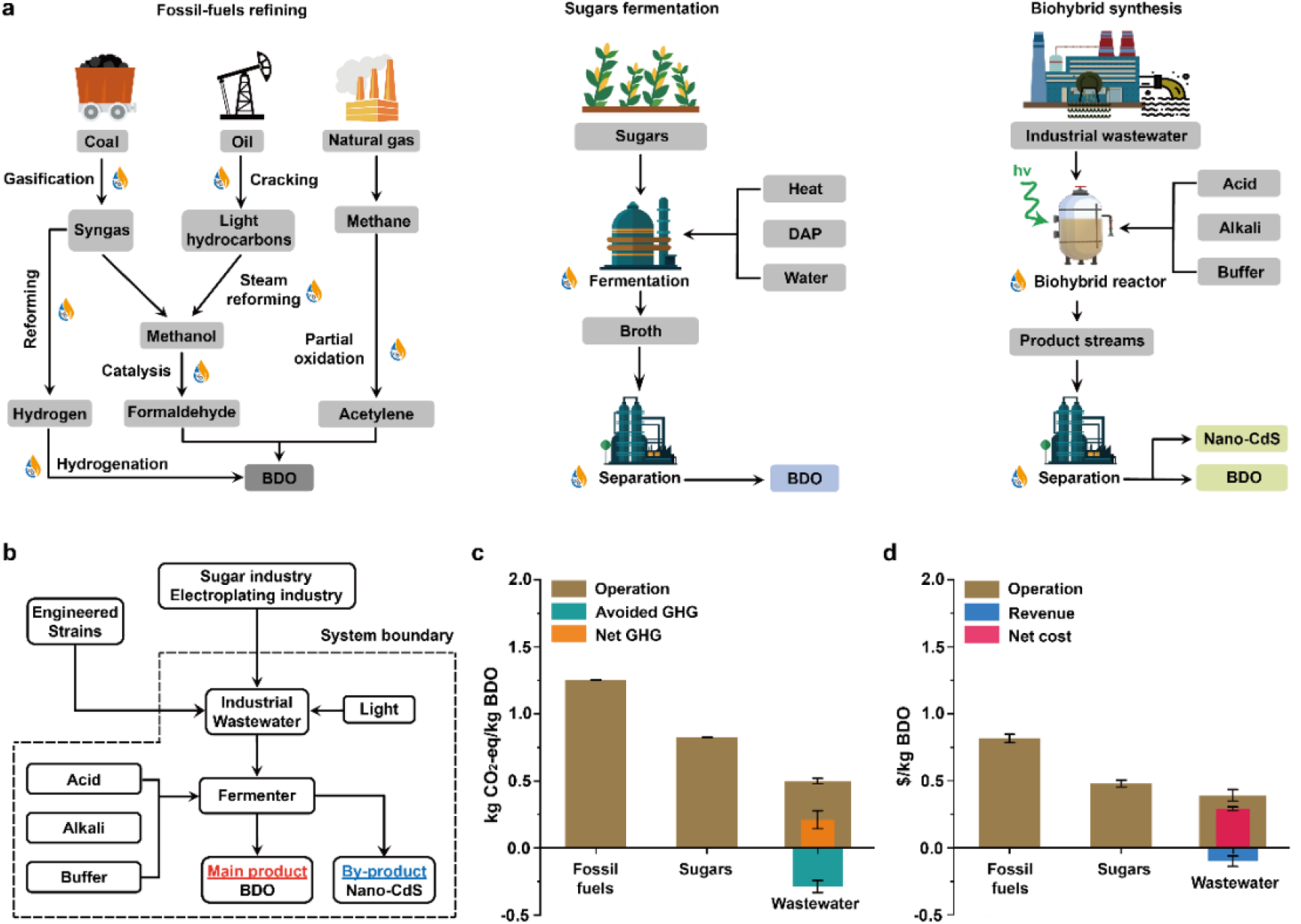
Life cycle analysis. **a)** Comparison of current routes to BDO by fossil-fuels refining, sugars fermentation, and solar-driven biohybrid synthesis from wastewater. Labelled steps represent energy consumption. **b)** System boundary of LCA for solar-driven biohybrid synthesis from wastewater, and its comparison in **c)** greenhouse gases (GHG) emissions and **d)** economic cost with current routes. Avoided treatment of industrial wastewater and by-products of nano-CdS offset partial GHG emissions and cost. Net GHG emission (orange) of 2,3-butanediol accumulates the positive emissions from operation (brown) of solar-driven biohybrid synthesis and avoided emissions (green) from industrial wastewater treatment, net cost (red) benefits from the revenues (blue) of avoided wastewater treatment and valuable by-products.

## Conclusion

Using wastewater resources as feedstock offers a cost-competitive and environmentally friendly approach for biomanufacturing. A tailored and engineered strain in this study is demonstrated to utilize multi-components in industrial wastewater to produce semiconductor biohybrids *in-situ* for solar-to-chemical production. The strain can consume organics for cell growth and recycle heavy metal ions and sulfate simultaneously into functional semiconductor for biohybrids production. Under light condition, semiconductors absorb light to generate photo-induced electrons for increasing NADH levels, stimulating central metabolism and enhancing BDO production yield in the biohybrids. Photo-bioreactor and LCA showed this method is potential able to scale up due to lower cost of feedstock and less CO_2_ emission, integrating synthetic biology and materials science and providing an alternative green future of sustainable biomanufacturing and environmental remediation.

## Supplementary Methods

1. Industrial wastewater.
2. Bacterial culture medium.
3. Engineering *Vibrio natriegens*.
4. Quantifying H_2_S gas production.
5. Biohybrids construction and heavy metals removal.
6. Scanning electron microscope (SEM).
7. Nanoparticles’ extraction.
8. Special aberration corrected transmission electron microscope (AC-TEM).
9. X-ray photoelectron spectroscopy (XPS).
10. X-ray powder diffraction (XRD).
11. Photoelectrochemical analysis.
12. Semiconductor biohybrid production from wastewater.
13. Preparation of semiconductor biohybrid for solar-driven BDO production.
14. Quantifying intracellular metabolites in semiconductor biohybrid system.
15. Quantifying BDO, acetoin and glucose.
16. *In-situ* production of semiconductor biohybrid to produce BDO using wastewater.
17. Scale-up production of semiconductor biohybrid to produce BDO using wastewater.
18. Life cycle analysis (LCA).

## Supplementary Information for LCA

1. Route 1. Production of 1,4-butanediol by fossil-fuels.
2. Route 2. Production of 2,3-butanediol by bio-fermentation.
3. Route 3. Production of 2,3-butanediol and CdS nanoparticles by solar-driven biohybrid from wastewater.
4. Potential offset of GHG emissions and economic cost.
5. Uncertainty analysis.

## Supplementary Methods

### Industrial wastewater

Three kinds of organic wastewater generated during production of biodiesel, sugar and corn starch, respectively, were obtained from factories in Harbin and Xingtai, China. These organic wastewaters have complex organic/inorganic compositions and known to contain major organic wastes of crude glycerol, molasses and starch, respectively, which are considered as pollutants in water since recycling of them by conventional physical/chemical approaches is uneconomical. We named these organic wastewaters as glycerol wastewater, molasses wastewater (Table S2, composition) and starch wastewater, respectively. Electroplating wastewater (Table S3, composition) was obtained from a metal electroplating factory in Shenzhen, China.

### Bacterial culture medium

**(1) Rich culture medium:** The LBv2 medium (per 1 L) contains LB powder 25 g, NaCl 11.9 g, KCl 0.313 g, and MgCl_2_ 2.2 g with corresponding antibiotics. All solidified media for bacterial growth contains 1.5% [w/v] agar. **(2) Well-defined minimum medium:** The modified M9 medium (per 1 L) contains NH_4_Cl 1 g, NaCl 12.4 g, Tris-HCl 10.99 g, Thiamine HCl 0.34 g, 40% glycerol 10 mL, casmino acid 1 g, MgSO_4_ 2 mM, CaSO_4_ 0.1 mM, KCl 0.313 g, MgCl_2_ 2.2 g, and pH was adjusted at 7.0 before use. **(3) Wastewater medium**: Wastewater medium was prepared by mixing electroplating wastewater and organic wastewater in a proportion and was amended with Cd^2+^, salts (per 1 L) of (NH_4_)_2_SO_4_ 5 g, NaCl 15 g, K_2_HPO_4_ 1 g, KH_2_PO_4_ 1 g, MgSO_4_ 0.25 g, CaCl_2_ 0.01 g in consideration of a tough condition of industrial wastewater with high salinity in practice. The main components and property of wastewater after pH adjustment were given in Table S4. *Vibrio natriegens* was routinely cultured using rich medium. The well-defined minimum medium or wastewater medium were used for the construction of biohybrids and chemical production, respectively.

### Engineering *Vibrio natriegens*

The bacterial strains are listed in Table S5, and the construction methods of engineering *V. natriegens* were described in detail.

#### Preparation of electrocompetent cells

*Escherichia coli* DH5α was routinely used for DNA manipulation. *V. natriegens* VnDx was used as starting strain in this study. *E. coli* was grown aerobically at 37 °C in Luria-Bertani (LB) broth or agar supplemented with 30 µg/ml chloramphenicol, 50 µg/ml kanamycin, or 100 µg/ml ampicillin, if required. *V. natriegens* was routinely grown at 37 °C in a rich culture medium with 30 µg/ml chloramphenicol, 200 µg/ml kanamycin, or 100 µg/ml ampicillin respectively if necessary. Electrocompetent cells were prepared in a manner similar to a previously described protocol^47^. Briefly, *V. natriegens* was grown in a flask in BHIv2 medium (37 g/L Teknova Brain Heart Infusion Broth Dry Media supplemented with additional salts of 204 mM NaCl, 4.2 mM KCl, and 23.14 mM MgCl_2_) overnight at 37 °C. This culture was inoculated (1:200 dilution) into BHIv2 culture and grown at 37 °C in a baffled flask to an OD_600_ of 0.5. The bacterial culture was incubated on ice for 30 min, and the pellets was collected and washed three times in ice-cold electroporation buffer (680 mM sucrose, 7 mM K_2_HPO_4_, pH 7). The electrocompetent cells was generated, and stored at -80 °C until use.

#### Electroporation

Electroporation protocol was adapted from previously described protocol^47^. Plasmid DNA (200 ng) and electrocompetent cells (100 μL) were gently mixed in a chilled 1.5-mL microcentrifuge tube. The cell–DNA suspension was transferred to a chilled electroporation cuvette with a 0.2-cm gap size. Cells were electroporated with 1.7 kV and immediately recovered in 500 µL recovery medium (BHI supplemented with v2 salts (204 mM NaCl, 4.2 mM KCl, 23.14 mM MgCl_2_, and 680 mM sucrose). The cells were incubated at 37 °C for 1 h and then plated on agar plates with appropriate antibiotic. The plates were incubated overnight at 37 °C for colonies growth.

#### Plasmids and strains construction

The DNA fragments contain cysteine desulfhydrase (*cdsH*) gene from *T. denticolawas* and the mutant serine acetyltransferase (*cysE*) gene with the promoter^27^ was chemically synthesized by Genscript. The synthetic DNA fragment was cloned into the plasmid vector pUC-GW-Kan to construct plasmid pXG203. The pXG203 was introduced into *V. natriegens* VnDx, resulting strain XG203. The BDO gene cluster containing its entire operon and the LysR-type transcriptional regulator from *Enterobacter cloacae subsp. dissolvens* SDM was amplified through PCR with the primer pair RABC-overlap-F (5’-3’: gcataatgcttaagtCTTCCTTTCGGGCTTTGTTAG) and RABC-overlap-R (5’-3’: tattgctcagcggtgGATATAGGCGCCAGCAAC). The linearized pACYCduet-1 vector was amplified through PCR with the primer pair pACYC-F (5’-3’: caccgctgagcaataactagc) and pACYC-R (5’-3’: acttaagcattatgcggccgc). The two fragments were assembled by Gibson Assembly, resulting in the plasmid pACYC-RABC. The plasmid pACYC-RABC was introduced into *V. natriegens* XG203 to generate strain XG203A.

### Quantifying H_2_S gas production

Quantitative H_2_S gas was monitored using IFU Hydrogen Sulfide Dräger-Tube^®^ (Dräger, Germany). The wild type *V. natriegens* and strain XG203 was cultured in a well-defined minimum medium at 37 °C for 72 hours with an initial OD_600_ of ∼0.6. A made-to-measure rubber stopper fitted by Dräger-Tube® were corked in a 250 ml Erlenmeyer flasks containing 50 ml cultures^17^. To quantify H_2_S gas production, the numbers on the Dräger-Tube® were recorded at specific time points.

### Biohybrids construction and heavy metals removal

*V. natriegens* XG203 was cultured in a rich culture medium at 37 °C for ∼3 hours, and then 1% bacterial culture was transferred into a fresh rich culture medium and grown at 37 °C for another ∼3 hours. Cells were collected after centrifugation and resuspended in a well-defined minimum medium with addition of various metal ions (CdCl_2_, PbCl_2_, CuSO_4_, ZnCl_2_, HgCl_2_, Na_2_SeO_3_, Bi(NO_3_)_3_·5H_2_O and HgCl_2_) separately, with an initial OD_600_ of ∼0.6. The concentration of all metal ions were 0.1 mM, but the Hg^2+^ was 0.01 mM. The bacterial cultures were grown at 37 °C for 24 hours. The supernatants were collected to evaluate the removal efficiency by inductively coupled plasma mass spectrometry (ICP-MS) (NexION 1000, PerkinElmer, USA). The pellets from bacterial culture with CdCl_2_ (0.1 mM) / PbCl_2_ (0.1 mM) / HgCl_2_ (0.01 mM) were collected to analyze biohybrids and nanoparticles, respectively.

### Scanning electron microscope (SEM)

To characterize semiconductor biohybrid, the pellets were collected and fixed by 2.5% glutaraldehyde and stored at 4 °C overnight. To remove the water content, the fixed pellets were sequentially submerged in a series of ethanol (30%, 50%, 70%, 80%, 90%, and 95%) for 15 min each and then in 100% ethanol for 20 min twice, ethanol: isoamyl acetate (v:v, 1:1) for 30 min, isoamyl acetate for one hour. A critical point dryer (Leica EM CPD30, Germany) was used to dry samples. The images were obtained using SEM (HITACHI, SU8010, Japan) with energy dispersive spectrometer (EDS) after gold spraying.

### Nanoparticles’ extraction

The pellets were collected and resuspended in Tris-HCl (50 mM, pH 7.5). Cell lysis was performed via sonication in an ice bath for 3 h with a probe sonicator (Scientz-IID, China). The samples were centrifuged at 13000 *g* for 1 h to collect the nanoparticles. To disperse nanoparticles, the particles were resuspended with Li_2_S/FA solution (5 g/L) through sonication for 15 min to obtain a uniform solution. Then the didodecyldimethylammonium bromide (DDAB)/toluene (50 g/L) was added to the top of the solution with continuous stirring. After the overnight reaction, the DDAB-modified nanoparticles were collected from the top layer of the solution (oil layer) and washed with acetone and ethanol three times to remove the excessive DDAB. The dispersion was then centrifuged at 10 000 rpm for 10 min and redispersed in toluene. The nanoparticles were collected for transmission electron microscopy, scanning transmission electron microscopy, and x-ray powder diffraction.

### Special aberration corrected transmission electron microscope (AC-TEM)

To observe the morphology of semiconductor biohybrid, the cross-sectional samples were prepared. The fixed pellets were centrifuged and washed by PBS (0.1 M, pH 7.4) for three times, and then suspended and wrapped in the 1% agarose solution before solidification. Agarose blocks were fixed with 1% OsO_4_ in PBS (0.1 M, pH 7.4) for 2 hours, and then rinsed in PBS (0.1 M, pH 7.4) for three times, 15 min each. For dehydration, the samples were exposed in a series of ethanol (30%, 50%, 70%, 80%, 90%, 95%, 100%, 100%) for 20 min and then submerged in acetone for 15 min twice.

The samples were exposed in acetone/EMBed 812 (1:1) for 2∼4 hours and acetone/EMBed 812 (1:2) overnight, and then in pure EMBed 812 for 5∼8 hours for resin penetration and embedding at 37 °C, and then inserted into the embedding models poured by the pure EMBed 812 at 37 °C overnight followed by polymerization at 65 °C for 48 hours in the oven. The resin blocks were cut to 60∼80 nm thin using the ultra microtome (UC7, Leica, Germany) by diamond knife (Ultra 45°, Daitome, Switzerland), and fished out onto the copper grid of 200 meshes. To observe the morphology of semiconductor biohybrid, the vaccum-dried copper grids was observed using AC-TEM (JEM-ARM300F, JEOL Ltd. Japan). To observe the morphology of nanoparticles, the dispersed suspension of nanoparticles were dropped onto copper grids and observed using AC-TEM (JEM-ARM300F, JEOL Ltd. Japan) after vaccum drying. High-angle annular dark-field (HAADF) STEM, energy dispersive spectrometer (EDS) mapping, and selective area electron diffraction (SAED) were performed at 300 kV.

### X-ray photoelectron spectroscopy (XPS)

To distinguish the valence state of element in semiconductor biohybrid, the freeze-dried pellets were analyzed by XPS (Nexsa, Thermo Scientific, USA) using Al Kα-ray at a work voltage of 15 kV using energy of 1486.6 eV, and the data were calibrated using C 1s at 284.80 eV.

### X-ray powder diffraction (XRD)

To figure out the crystal structure of semiconductor nanoparticles, XRD patterns were detected using XRD instrument (Areis, PANalytical, Netherlands) at 20∼80°.

### Photoelectrochemical analysis

The UV-Vis (Thermo Fisher Scientific, USA) was used to measure the direct bandgap. The electrochemical analysis was performed using Gamary instruments. The dispersed nanoparticles or semiconductor biohybrids were loaded on indium tin oxides (ITO) and carbon paper, respectively, to form a uniform 1 × 1 cm^2^ film by drop casting and vaccum drying. The photoelectrochemical measurements were performed by a standard three-electrode configuration in electrolyte (0.5 M Na_2_SO_4_). Ag/AgCl (3 M NaCl) and Pt wire served as the reference and counter electrode, respectively. A 300 W Xe-arc lamp (Newport, USA) equipped with a filter to block the infrared irradiation was employed as the solar simulator. A light meter with optothermal detector (THORLABS, PM400) was utilized to calibrate the light intensity to 1 sun illumination (100 mW cm^-2^). Current densities were recorded under 0.5 V bias vs Ag/AgCl.

### Semiconductor biohybrid production from wastewater

To demonstrate the capability to utilize organics in wastewater, *V. natriegens* XG203 was cultured using various wastewater-relevant organics (5 g/L) as carbon substrate in a well-defined minimum medium with addition of CdCl_2_ (0.1 mM) at 37 °C for 24 hours, with an initial OD_600_ of ∼0.6. To clarify the capability to utilize different sources of wastewater for semiconductor biohybrid production, *V. natriegens* XG203 was cultured in a well-defined minimum medium with addition of CdCl_2_ (0.01∼0.2 mM) at 37 °C for 24 hours, with an initial OD_600_ of ∼0.6. To explore the capability to produce semiconductor biohybrid using wastewater, strain XG203 was cultured in wastewater medium with organics supply by glycerol wastewater, molasses wastewater, and starch wastewater at 37 °C for 24 hours, with an initial OD_600_ of ∼0.6.

### Preparation of semiconductor biohybrid for solar-driven BDO production

To prepare semiconductor biohybrid, *V. natriegens* XG203A were inoculated into a rich culture medium to an OD_600_ of ∼1.0 and then transferred into a required large volume of rich culture medium with a 1% dilution. Cell pellets were collected at an OD_600_ of ∼1.0 and cultured in a well-defined minimum medium with addition of CdCl_2_ (0.1 mM) for the growth of *V. natriegens*-semiconductor hybrid at 37 °C, with an initial OD_600_ of ∼0.6. Semiconductor biohybrid were collected and cultured in a well-defined minimum medium with addition of glucose (5 g/L), cystein (1 mM) and flavine mononucleotide sodium (5 μM) at 37 °C under dark or light conditions (4.2 mW/cm^2^) for 24 hours. The light intensity was calibrated using a light meter with optothermal detector (THORLABS, PM400, USA). The supernatant was used to quantify BDO concentration, and the cell pellets were collected to quantify intracellular metabolites. The same condition was also used to culture bacterial system alone (without CdCl_2_) for BDO production under dark and light conditions as a control.

### Quantifying intracellular metabolites in semiconductor biohybrid system

To quantify intracellular metabolites, the extraction method of intracellular metabolites was according to that reported by Gao et al^48^. Briefly, cells were collected by centrifugation at 8000 *g* for 5 min at 4 °C and resuspended immediately in 2 mL of methanol/water (80:20, vol/vol) pre-cooled at -80 °C. The samples were centrifuged at 14 000 *g* for 10 min at 4 °C after incubation at -20 °C for 20 min. The supernatant was collected to quantify metabolites using liquid chromatography-tandem mass spectrometry (LC-MS/MS, SCIEX Triple Quad(tm) 5000+ QTRAP^®^ Ready, AB SCIEX, USA).

### Quantifying BDO, acetoin and glucose

To determine BDO concentration, bacterial cultures were centrifuged at 15000 rpm for 10 min and the supernatant was mixed with the isometric ethyl acetate with ultrasonic for 30 min and centrifugation for 10 min. Supernatant was filtered using nylon membrane (0.22 μm) and then determined using gas chromatography (GC). The GC system (Agilent 8890, Agilent Technologies Inc., USA) is equipped with a DB-WAX capillary column (30 m × 0.53 mm × 1 μm) and a flame ionization detector. Hydrogen gas was used as the carrier gas. The injector and detector temperature were maintained at 250 °C, and the oven temperature was 80 °C. The injection volume was 1 μL. To obtain the calibration curves for quantifying BDO and acetoin, a series of standards were quantified by GC system after the same pre-treatment. Glucose content was determined using glucose assay reagent (Beyotime, China).

### *In-situ* production of semiconductor biohybrid to produce BDO using wastewater

To demonstrate the capability of *in-situ* production of semiconductor biohybrid for BDO production using wastewater, *V. natriegens* XG203A was inoculated into a rich culture medium and cultured at 37 °C until OD_600_ ∼1.0 and then transferred into a fresh rich culture medium with 1% dilution. When an OD_600_ of ∼0.6 was reached, the concentrated cultures were resuspended and cultured using molasses wastewater (containing 4.68 g/L sugar) amended with salts (referred in the wastewater medium) at 37 °C until OD_600_ ∼1.0. Cell pellets were collected and cultured using wastewater medium with organics (4.10 g/L sugar) supply by molasses wastewater at 37 °C for 1 h under dark and light conditions (4.2 mW/cm^2^) with an initial bacterial OD_600_ of ∼1.0. Cystein (1 mM) and flavine mononucleotide sodium (5 μM) were added in the cultures. The same condition was also used to culture bacterial system alone in molasses wastewater amended with salts (referred in the wastewater medium) for BDO production under dark and light conditions as the control. The light intensity was calibrated using a light meter with optothermal detector (THORLABS, PM400, USA). The supernatant was used to detect BDO, acetoin, and total sugar (anthrone-sulfuric acid method). To explore the capability to utilize wastewater containing high concentration of molasses (3.0∼48.0 g/L) to produce semiconductor biohybrid for BDO production, *V. natriegens* XG203A was also cultured in wastewater medium with organics supply by different concentrations of molasses wastewater at 37 °C for 24 hours under light condition (4.2 mW/cm^2^) with an initial bacterial OD_600_ of ∼1.0.

### Scale-up production of semiconductor biohybrid to produce BDO using wastewater

To scale up *in-situ* production of semiconductor biohybrid and BDO, *V. natriegens* XG203A was cultured using wastewater medium in a 5-L fermenter (Biostat® A, Sartorius Stedim Biotech, Germany). Briefly, when an OD_600_ of XG203A cultured in a rich culture medium reached ∼1.0, the bacterial cultures were transferred into 4 L of molasses wastewater (containing 19.2 g sugar) amended with salts (referred in the wastewater medium) by a 2% dilution. When an OD_600_ was reached at ∼1.0, the concentrated cultures were transferred into a 5-L fermenter containing 3 L of wastewater medium with organics supply by molasses wastewater (containing 52.2 g sugar) under an illumination of 6 mW/cm^2^. The parameters of fermenter were set at 37 °C, pH 7.0, 600 rpm, and an air flow of 500 ccm. The samples were collected at a specific time interval to detect BDO, acetoin, and total sugar.

### Life cycle analysis (LCA)

As the impacts associated with construction of facilities, transportation of products, and treatment of waste streams are not considered in the system boundary of renewable fuel standard, LCA is conducted to analyze the greenhouse gases (GHG) emissions and economic cost with the scope of cradle-to-gate excluding the utilities constructure and end-of-life disposal^49,50^. Life-cycle inventory and emission factors in the production of 1 kg BDO using conventional routes of fossil-fuels refining and bio-fermentation of pure sugars are calculated using Ecoinvent database and models from Argonne National Laboratory^49,51^. A modified industrial model of bio-chemicals fermentation is adopted here to comprehensively indicate the environmental and economic impacts of BDO production with less uncertainties according to the lab-data of solar-driven biohybrid synthesis using wastewater. Conventional treatments of industrial wastewater consume chemicals and energy with auxiliary GHG emissions and investment. LCA was used to analyze the potential offset of GHG emission and economic cost by the solar-driven biohybrid synthesis via system expansion, wherein the wastewater treatment is avoided and valuable by-products (nano-CdS) are produced. Life-cycle inventory and parameters in the detailed calculations are provided in the Supplementary Information for LCA, including consumption of materials and energy for BDO production and conventional wastewater treatments. IPCC-2021-GWP100^52^ is conducted to quantify the GHG emission. An extended life-cycle economic evaluation based on the similar goal and scope of LCA was used to calculate the economic burdens of the whole life-cycle process^53^.

### Supplementary Information for LCA

#### Route 1. Production of 1,4-butanediol by fossil-fuels

Hydrogenation of butyne-diol (the intermediate of Reppe process with feedstocks of formaldehyde and gaseous acetylene) is often used in the industrial production of 1,4-butanediol by fossil-fuels^54^. Formaldehyde derives from methanol production by the steam reforming of light hydrocarbons and syngas, which is catalyzed by metal-oxide or sliver particles in 600-700 °C^55^. The off-gas from methanol conversion can be reutilized to thermally activate the spent catalysts, thus no consumption of catalysts for formaldehyde production is assumed. The partial oxidation of methane produces acetylene streams with a certain amount of H_2_ and CO, which can be further separated as syngas for hydrogenation of butyne-diol^56^. Approximate one third of the methane in natural gas converts to acetylene while the rest is combusted to thermal steam^57^. The residual heat of steam is assumed to be recovered in ∼70% for heating of reactors. Besides GHG emissions from reactors operation and chemicals consumption, partial oxidation of methane generates additional GHG emissions (0.12 kg CO_2_-eq/kg acetylene by mass allocation) in the acetylene (∼16%, wt) streams^56^. Methane in the acetylene streams is assumed to be recovered without contribution to GHG emission. Fig. S6 exhibits the schematic of 1,4-butanediol production. Table S6 summarizes the energy and chemicals consumed in the production of 1,4-butanediol, and Fig. S7 demonstrates the detailed contributions of factors to GHG emission and cost.

#### Route 2. Production of 2,3-butanediol by bio-fermentation

2,3-butanediol is the major replacement of 1,4-butanediol due to their isomerism, similar properties, and main sources from bio-fermentation of sugars (sucrose, fructose, glucose, etc.). The slurry of sugars, as feedstock^58^, is converted to 2,3-butanediol via feedstock sterilization, fermentation, and products separation^59,60^. The accessory fermentation ingredient, corn steep liquor (CSL), was mixed with the slurry of sugars and heated at 121 °C for ∼15 min with natural cooling. Bio-fermentation is conducted at 37 °C with residence of 48 hours, wherein CSL and added diammonium phosphate (DAP) serve as the nitrogen sources. The fermentation broth is then flashed to purge the gases, centrifuged to separate the cell-mass from liquid streams, and transferred to the separation unit. The low concentration and high solubility hinder the separation of 2,3-butanediol from fermentation system, reducing the economic benefits of conventional multi-stage distillation^61^. A hybrid extraction-distillation was proposed here with solvents of 1-butanol due to its favorable selectivity and relatively low boiling-point^62^. Although other technologies like membrane separation and salting-out extraction are potentially feasible, the solvent extraction has been identified as industrially promising in the long term^63^. The mass-flow rate of solvent transferred to the multi-stage mixers is assumed to be 1.2 times than the water in product streams of fermentation broth. The extract solvent is distilled at 0.06 atm and ∼120 °C with lower heating requirement than conventional distillation (∼252 °C), while residues of 2,3-butanediol and water are then sequentially distilled at 0.2 atm by conventional distillation. Fig. S8 exhibits the schematic of 2,3-butanediol production. Table S7 summarizes the energy and materials consumed in the production of 2,3-butanediol, and Fig. S9 demonstrates the detailed contributions of factors to GHG emission and cost.

#### Route 3. Production of 2,3-butanediol and CdS nanoparticles by solar-driven biohybrid from wastewater

To make the comparison of environmental and economic impacts of different routes of 2,3-butanediol production credible, similar industrial fermentation of biochemicals is adopted here as the industrial modelling to investigate the potential advantages of solar-driven biohybrid synthesis, which also complies with the standards of LCA processes due to uncertainties in the lab-scale processes and incomprehensiveness in indicating the related impacts. Thus, lab-scale production of 2,3-butanediol by solar-driven biohybrid using wastewater is assumed to be modified with an industrial modelling similar to the described routes of bio-fermentation, which includes the culture of *V. natriegens* from mature seed using wastewater medium in an irradiated fermenter, conditioning of pH by hydrochloric acid and caustic soda with buffer solution, and separation of target products and by-products. Fig. S10 exhibits the schematic of 2,3-butanediol production by the solar-driven biohybrid. LCA with scope of cradle-to-gate is analyzed for **Route 3**, and similar strategy for the separation of 2,3-butanediol by the hybrid extraction-distillation is assumed^63^. Besides 2,3-butanediol as target products, *in-situ* produced nano-CdS as significant semiconductor materials are also assumed to be recovered via ultrasound cell-lysis and centrifugation after per cycle of operation. Table S8 summarizes the energy and materials consumed in the production of 2,3-butanediol and by-products of nano-CdS according to the modified industrial modelling of 2,3-butanediol production by bio-fermentation, and Fig. S11 demonstrates detailed contributions of factors to GHG emission and cost.

### Potential offset of GHG emissions and economic cost

The strategy of solar-driven biohybrid synthesis from wastewater produces valuable products and avoids the cost and potential GHG emissions from wastewater treatment^64^. Meanwhile, valuable by-products of nano-CdS can offset partial cost of 2,3-butanediol production. LCA analyzed above eco-friendly and economic advantages of solar-driven biohybrid synthesis via system expansion eliminating wastewater treatment^16^. Typical wastewater treatment process, anaerobic-anoxic-aerobic (A^2^O) as the secondary treatment with front-end primary treatment, is assumed here to treat the molasses wastewater with high COD^53,65^, while coagulation/flocculation-sedimentation-filtration-ultraviolet reduction as the tertiary treatment is assumed here to concentrate heavy-metal ions in electroplating wastewater, and functional unit of wastewater treatment is 1 m^3^ influent^66^. The residual sludge from wastewater treatment is thickened, dewatered, and transferred to the landfill sites^53^. Wastewater treatment is a particularly complex system, but the environmental and economic burdens of construction and demolition stages of utilities are negligible compared to the operation stages, wherein the consumption of electricity and chemicals contribute up to ∼90% of environmental impacts^67-69^. Therefore, this scenario only considered the operation stages of wastewater treatment with system boundary shown in Fig. S12. Table S9 summarizes the consumption of energy and materials of wastewater treatment, emission factors of CH_4_ and N_2_O from A^2^O operation are provided in Table S10^70^. Periodic market prices of energy and chemicals consumed, and by-products are summarized in Table S11^53^. Estimated unit offset of GHG emission and economic cost are calculated in equation S1-S3, results are summarized in Table S12 and exhibited in Fig. 5c and 5d.

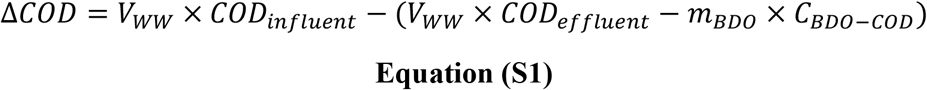

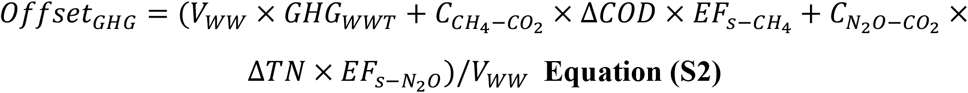

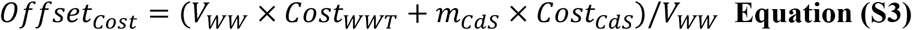

where ***COD***_***influent***_ and ***COD***_***effluent***_ are the concentrations of COD (kg/m^3^) in the influent and effluent of industrial wastewater, ***m***_***BDO***_ is the mass (kg) of produced BDO in one cycle of operation, ***C***_***BDO−COD***_(∼1.95 kg/kg) is the conversion constant of COD to per kilogram of BDO, ***offset***_***GHG***_ (kg CO_2_-eq/m^3^) is the offset of GHG (CO_2_, CH_4_, and N_2_O) emission from wastewater treatment, ***V***_***WW***_ (m^3^) is the treated volume of wastewater, ***GHG***_***WWT***_ (kg CO_2_-eq/m^3^) is the GHG emission of wastewater treatment, **Δ*COD***(kg) is the removal amount of COD before and after wastewater treatment, **Δ*TN*** (kg) is the removal amount of total-nitrogen before and after wastewater treatment, 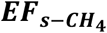 (10^−3^ kg CH_4_/kg COD removal) and 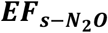 (10^−3^ kg N_2_O/kg TN removal) are the emission factors of CH_4_ (generated from anaerobic zone) and N_2_O (generated from aerobic zone) during A^2^O treatment, 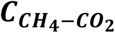 (∼25, dimensionless) and 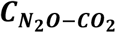 (∼298, dimensionless) are the constant of GHG effects of CH_4_ and N_2_O compared against CO_2_ according to the reports of Intergovernmental Panel on Climate Change, ***offset***_***Cost***_ ($/m^3^) is the offset of economic burdens from wastewater treatment and revenues from valuable by-products of nano-CdS, ***Cost***_***WWT***_ ($/m^3^) is the economic expense of wastewater treatment, ***m***_***CdS***_ (kg) is the production mass (dry-weight) of nano-CdS by solar-driven biohybrid synthesis, ***Cost***_***CdS***_ ($/kg) is the market prices of nano-CdS in purity of semiconductor class.

### Uncertainty analysis

To characterize the uncertainty of GHG emissions, economic cost of BDO and CdS nanoparticles by different routes, Monte Carlo simulation with Latin hypercube sampling (10000 trials) was used to propagate input uncertainty of 49 parameters (Table S13) to quantify GHG emissions and costs.

## Acknowledgements

This work was supported by the National Natural Science Foundation of China (Grant No. 32230060, C. Y. and X.G.; Grant No. 22176046, L.L.; Grant No. 32171426, X.G.; Grant No. 52200090, S.P.), Shenzhen Science and Technology Program (Grant No. GXWD20220811173949005, KQTD20190929172630447 and JCYJ20210324124209025, L.L.; Grant No. JCYJ20220818101804010, X.G.), the National Key R&D Program of China (Grant No. 2021YFA0910800, X.G.), State Key Laboratory of Urban Water Resource and Environment (Harbin Institute of Technology) (Grant No. 2021TS13, L.L.), and Natural Science Foundation of Guangdong Province (Grant No. 2022A1515012016, L.L.).

## Author Contribution

X. G., L. L., Y. L., and C. Y. supervised the research. X. G. and L. L. designed the experiments. S. P., W. Y., and W. F. contributed to the biohybrids production, structural and chemical characterizations. W. C., and S. P. performed the metabolic experiment, with the results verified by X. G. S. P., R. Y., and W. Y. contributed to the wastewater-relevant experiments and fermenter data. L. C., Z. L., R. Y., and W. C. performed the photoelectrochemical analysis. W. C., X. G., and L. L. contributed to the LCA data. S. P., X. G., Y. L., L. L., W. Y., and W. C. wrote the manuscript, and received comments from all the other authors.

## Declaration of Interests

L. L., X. G., R. Y., S. P., and W. Y. are co-inventors on filed China patents CN202310145122.9 and CN202210318999.9 related to produce semiconductor nanoparticles and biohybrids directly from wastewater by engineered strains that incorporates discoveries incorporated in this manuscript.

