## Supplementary Information 1 for "Scalable Solar-Driven Chemical Production by Semiconductor Biohybrids Synthesized from Wastewater Pollutants"

### Supplementary Tables

**Table S1.** Comparison of the residual metal concentration (μg/L) with sewage effluent standard.

**Table S2.** Compositions of molasses wastewater used in this study.

**Table S3.** Compositions of the electroplating wastewater.

**Table S4.** Main characteristics of wastewater for BDO production in flask and scale-up bioreactor culture.

**Table S5.** Bacterial strains and plasmids used in this study.

**Table S6.** Energy and material inventory of 1,4-butanediol production by fossil-fuels refining.

**Table S7.** Energy and material inventory of 2,3-butanediol production by bio-fermentation.

**Table S8.** Energy and material inventory of 2,3-butanediol production by solar-driven biohybrid using wastewater.

**Table S9.** Energy and material inventory of conventional industrial wastewater treatment.

**Table S10.** Emission factors (EFs) of GHG ($CH_4$ and $N_2O$) of wastewater treatment by $A^2O$ technology.

**Table S11.** Market prices of electricity and chemicals consumed in wastewater treatment and nano-CdS.

**Table S12.** Estimated unit offset of GHG emission and economic cost from wastewater treatment and valuable by-products, nano-CdS.

**Table S13.** Parameters and probability distributions used in the illustrative routes.

### Supplementary Figures

**Figure S1** Other evidences for the production of CdS semiconductor biohybrid.

**Figure S2** Bacterial capability for the production of various semiconductor biohybrids using different heavy metals.

**Figure S3** Production of PbS semiconductor biohybrid.

**Figure S4** Production of HgS semiconductor biohybrid.

**Figure S5** Solar-driven production of acetoin, a direct precursor for BDO production.

**Figure S6** Schematic and system boundary of 1,4-butanediol production by fossil-fuels refining.

**Figure S7** Contributions of factors to GHG emission and cost of 1,4-butanediol production by fossil-fuels refining.

**Figure S8.** Schematic and system boundary of 2,3-butanediol production by fermentation of sugars.

**Figure S9** Contributions of factors to GHG emission and cost of 1,4-butanediol production by fermentation of sugars.

**Figure S10** Schematic and system boundary of 2,3-butanediol production by solar-driven biohybrid using wastewater.

**Figure S11** Contributions of factors to GHG emission and cost of 2,3-butanediol production by solar-driven biohybrid using wastewater.

**Figure S12** Schematic and system boundary of conventional wastewater treatment process used for treating industrial wastewater with high COD and heavy metals.

### Supplementary Tables

**Supplementary Table S1 | Comparison of the residual metal concentration with sewage effluent standard.**

| Heavy metals | Concentration (μg/L) | GB/T 31962-2015 (China) | EPA (USA)* |
|---|---|---|---|
| **Cd** | 7.10 | 50 | 17.2~1200 |
| **Pb** | 9.50 | 500 | 57~3400 |
| **Cu** | 41.42 | 2000 | 23~5000 |
| **Hg** | 1.17 | 5 | 0.0018~110 |
| **Se** | 80.72 | 500 | 5~1600 |
| **Bi** | 7.06 | / | / |
| **Zn** | 2678.28 | 5000 | 82~114000 |

**Note**: *Limitations are daily maximum of pollutant limitations in the industrial wastewater according to Effluent Guidelines Database. The residual concentrations of heavy metal in the liquid supernatant were detected after eliminating the pellets of strain XG203 cultured in a well-defined minimum medium with addition of one kind of heavy metals. The concentration of all metal ions were 0.1 mM, but the $Hg^{2+}$ was 0.01 mM. The residual concentrations of each heavy metal are lower than that of the limitation by GB/T 31962-2015 (China) and EPA (USA), except that the effluent limit of bismuth is not referred in the standard.

**Supplementary Table S2 | Compositions of molasses wastewater used in this study.**

| pH | Conductivity (μS/cm) | TOC (mg/L) | COD (mg/L) | $BOD_5$ (mg/L) | TN (mg/L) | Color | Viscosity (mPa×S) |
|---|---|---|---|---|---|---|---|
| 4.12 | 6830 | $8.78\times10^3$ | $2.54\times10^4$ | $4.51\times10^3$ | 826 | $5\times10^3$ | 3.5 |
| **TS (g/L)** | **Sucrose (g/L)** | **Fructose (g/L)** | **Glucose (g/L)** | **$SO_4^{2-}$ (g/L)** | **$NO_3^-$ (g/L)** | **$NO_2^-$ (g/L)** | **$NH_4^+$ (g/L)** |
| 41.5 | 20.8 | 8.1 | 2.0 | 1.6 | 0.1 | 0.2 | 0.1 |
| **K** | **Mg** | **Ca** | **Fe** | **Na** | **Ti** | **Zn** | **Mn** |
| 9.44 | 3.068 | 2.38 | 0.179 | 0.05 | 0.017 | 0.011 | 0.011 |

**Note**: The indicators (pH, conductivity, TOC, COD, $BOD_5$, TN, color, and viscosity) were detected from wastewater containing 24 g/L of molasses, while other compositions were detected from raw molasses wastewater. TOC, total organic carbon; COD, chemical oxygen demand; $BOD_5$, biochemical oxygen demand; TN, total nitrogen; TS, total sugars; Sucrose, fructose and glucose were detected using high performance liquid chromatography; $SO_4^{2-}$, $NO_3^-$, $NO_2^-$, and $NH_4^+$ were detected using ion chromatography; Metal ions (g/kg) were detected using ICP.

**Supplementary Table S3 | Compositions of the electroplating wastewater.**

| pH | Conductivity (μS/cm) | TOC (mg/L) | $NH_4^+$ (mg/L) | $NO_3^-$ (mg/L) | $NO_2^-$ (μg/L) | TN (mg/L) | $SO_4^{2-}$ (mg/L) |
|---|---|---|---|---|---|---|---|
| 3.51 | 2660 | 40 | 20 | 0.1327 | 0.036 | 5.00 | 5.97 |
| **Ni** | **Cd** | **Pb** | **Hg** | **Mg** | **Cr** | **Fe** | **Al** |
| 15.5 | 22.08 | 19.85 | 18.99 | 2.46 | 6.18 | 5.77 | 6.47 |

**Note**: Metal ions (mg/L) were detected using ICP.

**Supplementary Table S4 | Main characteristics of wastewater for BDO production in flask and scale-up bioreactor culture.**

| Scale | pH | TS (mg/L) | COD (mg/L) | $SO_4^{2-}$ (mg/L) | TN (mg/L) | Cd (mM) |
|---|---|---|---|---|---|---|
| **flask** | 7.0 | $4.1\times10^3$ | $1.0\times10^4$ | $4.02\times10^3$ | $2.5\times10^3$ | 0.1 |
| **bioreactor** | 7.0 | $52.2\times10^3$ | $9.4\times10^4$ | $8.74\times10^3$ | $4.5\times10^3$ | 0.1 |

**Note**: Cd content was detected using ICP-MS.

| Supplementary Table S5 丨 Bacterial strains and plasmids used in this study. | | |
|---|---|---|
| **Strains or plasmids** | **Description** | **References** |
| **Bacterial strains** | | |
| *Vibrio natriegens* VnDx | Derived from ATCC 14048 by integrating the T7 RNA polymerase expression cassette at the dns locus | [1] |
| *Escherichia coli* DH5α | | Novagen |
| *V. natriegens* XG203 | VnDx harboring pXG203 | This work |
| *V. natriegens* XG203A | XG203 harboring pACYC-RABC | This work |
| **Plasmids** | | |
| pET-RABC | Used to offer the BDO gene cluster containing its entire operon and the LysR-type transcriptional regulator from *Enterobacter cloacae subsp. dissolvens* SDM | [2] |
| pUC-GW | Expression vector, $Kan^r$ | GENEWIZ |
| pXG203 | pUC-GW containing the designed DNA fragment that the recA (ΔLexA) promoter fused with the cysteine desulfhydrase (*cdsH*) gene from *T. denticolawas* and the mutant serine acetyltransferase (*cysE*) gene | This work |
| pACYCduet-1 | Expression vector, $Cm^r$ | Novagen |
| pACYC-RABC | pACYCduet-1 containing the BDO gene cluster containing its entire operon and the LysR-type transcriptional regulator from *Enterobacter cloacae subsp. dissolvens* SDM | This work |

**Supplementary Table S6 | Energy and material inventory of 1,4-butanediol production by fossil-fuels refining.**

| | Inputs | Usage amount | Units | Description in database |
|---|---|---|---|---|
| **Production of Acetylene** | Natural gas | 0.65 | $Nm^3/kg$ | Natural gas, high pressure {GLO}\| market group for \| Cut-off, S |
| | Electricity | 0.35 | kWh/kg | Electricity, high voltage {GLO}\| market group for \| Cut-off, S |
| | Oxygen | 0.82 | kg/kg | Oxygen, liquid {RoW}\| market for \| Cut-off, S |
| | N-methy-2-lpyrrolidone | 0.083 | kg/kg | N-methyl-2-pyrrolidone {GLO}\| market for \| Cut-off, S |
| **Production of formaldehyde** | Electricity | 0.17 | kWh/kg | Electricity, high voltage {GLO}\| market group for \| Cut-off, S |
| | Methanol | 1.20 | kg/kg | Methanol {GLO}\| market for \| Cut-off, S |
| **Production of 1,4-butanediol** | Natural gas | 0.020 | $Nm^3/kg$ | Natural gas, high pressure {GLO}\| market group for \| Cut-off, S |
| | Acetylene | 0.28 | kg/kg | —— |
| | Formaldehyde | 0.66 | kg/kg | —— |
| | Hydrogen | 0.050 | kg/kg | Hydrogen, gaseous {GLO}\| market for hydrogen, gaseous \| Cut-off, S |

**Note:** Life-cycle inventory of 1,4-butanediol production by fossil-fuel refining is derived from the model of Greenhouse gases, Regulated Emissions, and Energy use in Transportation (GREET), Argonne National Laboratory. Ecoinvent 3.8 embedded in Simapro 9.4 is the front-end database of inventory. IPCC-2021-GWP100 for the quantitative evaluation of GHG emission determines the GHG emission of 1,4-butanediol production. {GLO}, global; {RoW}, rest of world.

**Supplementary Table S7 | Energy and material inventory of 2,3-butanediol production by bio-fermentation.**

| Inputs | Usage amount | Units | Description in database |
|---|---|---|---|
| Natural gas | 0.30 | $Nm^3$/kg | Natural gas, high pressure {GLO}\| market group for \| Cut-off, S |
| Electricity | 2.2E-04 | kWh/kg | Electricity, high voltage {GLO}\| market group for \| Cut-off, S |
| Sugars | 0.70 | kg/kg | Sugar, from sugarcane {GLO}\| market for \| Cut-off, S |
| Diammonium phosphate (DAP) | 0.015 | kg/kg | Diammonium phosphate {RoW}\| market for diammonium phosphate \| Cut-off, S |
| Corn steep liquor (CSL) | 0.092 | kg/kg | Corn steep liquor/kg/RNA |
| 1-butanol | 0.028 | kg/kg | 1-butanol {GLO}\| market for \| Cut-off, S |
| Yeast | 7.6E-03 | kg/kg | Fodder yeast {RoW}\| ethanol production from whey \| Cut-off, S |

**Note:** Sugars in this inventory mainly contain sucrose, fructose, glucose, etc. with compositions provided in Table S2.

**Supplementary Table S8 | Energy and material inventory of 2,3-butanediol production by solar-driven biohybrid using wastewater.**

| Production of 2,3-butanediol and CdS nanoparticles | | | |
|---|---|---|---|
| **Inputs** | **Usage amount** | **Units** | **Description in database** |
| Natural gas | 0.30 | $Nm^3$/kg | Natural gas, high pressure {GLO}\| market group for \| Cut-off, S |
| Electricity | 6.27E-03 | kWh/kg | Electricity, high voltage {GLO}\| market group for \| Cut-off, S |
| Water | 80.66 | kg/kg | Water, deionised {RoW}\| water production, deionised \| Cut-off, S |
| 1-butanol | 0.028 | kg/kg | 1-butanol {GLO}\| market for \| Cut-off, S |
| Caustic soda | 0.095 | kg/kg | Hydrochloric acid, without water, in 30% solution state {RoW}\| market for \| Cut-off, S |
| Hydrochloric acid | 0.089 | kg/kg | Sodium hydroxide, without water, in 50% solution state {GLO}\| market for \| Cut-off, S |
| Buffer solution | 2.37 | L/kg | —— |
| Molasses waste | 10.25 | $m^3$/kg | —— |
| Electroplating wastewater | 2.85E-03 | $m^3$/kg | —— |

**Note:** Buffer solution contains $(NH_4)_2SO_4$ (10 ppm), NaCl (30 ppm), $K_2HPO_4$ (2 ppm), $MgSO_4$ (0.5 ppm), $CaCl_2$ (0.02 ppm) in neutrality modified by caustic acid, which functions as the replacement of DAP (nitrogen sources of microbials) in modeling of bio-fermentation. Life-cycle inventory in Table S8 corresponds to the functional unit of 1 kg 2,3-butanediol as the major product, while the yield of nano-CdS as by-products is determined through per cycle of 2,3-butanediol production.

**Supplementary Table S9｜Energy and material inventory of conventional industrial wastewater treatment.**

| Inputs | Usage amount | Units | Description in database | Outputs | Emission amount | Units | Description in database |
|---|---|---|---|---|---|---|---|
| **Primary treatment** | | | | | | | |
| TSS | 1.14E-01 | kg/m$^3$ | [a] | Grid slag | 6.58E-02 | kg/m$^3$ | Waste treatment, Landfill of waste, Slag/ash, EU27 |
| COD | 3.13E-01 | kg/m$^3$ | COD, Chemical Oxygen Demand, water, river | Sanding | 3.84E-02 | kg/m$^3$ | Waste foundry sand {GLO}\| market for waste foundry sand \| Cut-off, S |
| $BOD_5$ | 1.11E-01 | kg/m$^3$ | $BOD_5$, Biological Oxygen Demand, water, river | $H_2S$ | 2.60E-06 | kg/m$^3$ | Hydrogen sulfide {GLO}\| market for \| Cut-off, S |
| $NH_3$-N | 2.24E-02 | kg/m$^3$ | Ammonium, ion, water, river | $NH_3$ | 1.45E-05 | kg/m$^3$ | Ammonia, air, high population density |
| TP | 2.86E-03 | kg/m$^3$ | Phosphate, water, river | | | | |
| TN | 2.65E-02 | kg/m$^3$ | Ammonium, ion, water, river<br>Nitrite, water, river<br>Nitrate, water, river<br>Nitrogen, organic bound, water, river | | | | |
| Electricity | 5.32E-02 | kWh/m$^3$ | Electricity, high voltage {GLO}\| market group for \| Cut-off, S | | | | |
| **Secondary treatment** | | | | | | | |
| **Inputs** | **Usage amount** | **Units** | **Description in database** | **Outputs** | **Emission amount** | **Units** | **Description in database** |
| Acetic acid | 3.29E-03 | kg/m$^3$ | Acetic acid, without water, in 98% solution state {GLO}\| market for \| Cut-off, S | $H_2S$ | 5.53E-06 | kg/m$^3$ | Hydrogen sulfide {GLO}\| market for \| Cut-off, S |
| Electricity | 2.22E-01 | kg/m$^3$ | Electricity, high voltage {GLO}\| market group for \| Cut-off, S | $NH_3$ | 2.77E-05 | kg/m$^3$ | Ammonia, air, high population density |
| **Tertiary treatment** | | | | | | | |
| **Inputs** | **Usage amount** | **Units** | **Description in database** | **Outputs** | **Emission amount** | **Units** | **Description in database** |
| Electricity | 2.88E-01 | kWh/m$^3$ | Electricity, high voltage {GLO}\| market group for \| Cut-off, S | TSS | 7.00E-03 | kg/m$^3$ | [a] |
| Polyacrylamide | 3.00E-04 | kg/m$^3$ | Polyacrylamide {GLO}\| market for \| Cut-off, S | COD | 2.10E-02 | kg/m$^3$ | COD, Chemical Oxygen Demand, water, river |
| Lime | 7.00E-03 | kg/m$^3$ | Quicklime, milled, loose {RoW}\| market for quicklime, milled, loose \| Cut-off, S | $BOD_5$ | 8.00E-03 | kg/m$^3$ | $BOD_5$, Biological Oxygen Demand, water, river |
| Sand | 1.25E-03 | kg/m$^3$ | Sand {GLO}\| market for \| Cut-off, S | $NH_3$-N | 6.49E-04 | kg/m$^3$ | Ammonium, ion, water, river |
| Gravel | 7.20E-04 | kg/m$^3$ | Gravel, round {RoW}\| gravel and sand quarry operation \| Cut-off, S | TP | 8.70E-05 | kg/m$^3$ | Phosphate, water, river |
| | | | | TN | 7.96E-03 | kg/m$^3$ | Ammonium, ion, water, river<br>Nitrite, water, river<br>Nitrate, water, river<br>Nitrogen, organic bound, water, river |
| **Sludge treatment** | | | | | | | |
| **Inputs** | **Usage amount** | **Units** | **Description in database** | **Outputs** | **Emission amount** | **Units** | **Description in database** |
| Aluminum chloride | 8.22E-02 | kg/m$^3$ | Aluminium chloride {GLO}\| aluminium chloride production \| Cut-off, S | Biochemical sludge | 3.79E-01 | kg/m$^3$ | Raw sewage sludge (waste treatment) {GLO}\| market for \| Conseq, S |
| Polyacrylamide | 8.22E-04 | kg/m$^3$ | Polyacrylamide {GLO}\| market for \| Cut-off, S | $H_2S$ | 6.60E-07 | kg/m$^3$ | Hydrogen sulfide {GLO}\| market for \| Cut-off, S |
| Electricity | 5.60E-03 | kWh/m$^3$ | Electricity, high voltage {GLO}\| market group for \| Cut-off, S | $NH_3$ | 4.11E-06 | kg/m$^3$ | Ammonia, air, high population density |

**Note: a**: TSS represents the concentration of salt and heavy-metal ions in the effluent. The input items covered energy and chemical consumption in each treatment stage. The output items involved pollutants in the effluent, including biochemical oxygen demand ($BOD_5$), chemical oxygen demands (COD), ammonia nitrogen ($NH_3$-N), total nitrogen (TN), total phosphorus (TP), and total suspended solids (TSS), waste gas (ammonia ($NH_3$) and hydrogen sulfide ($H_2S$)), and solid waste (grid slag, sanding, biochemical sludge). Other GHG emissions ($CH_4$ and $N_2O$) besides indirect $CO_2$ emission from electricity consumption are not discussed here, and calculation is according to emission factors in Table S10. The scale of wastewater treatment is set as $1\times10^4$ m$^3$ per day.

**Supplementary Table S10 | Emission factors (EFs) of GHG ($CH_4$ and $N_2O$) of wastewater treatment by $A^2O$ technology.**

| EFs of $CH_4$ ($10^{-3}$ kg $CH_4$/kg COD removal) | Distribution | EFs of $N_2O$ ($10^{-3}$ kg $N_2O$/kg TN removal) | Distribution |
|---|---|---|---|
| 1.7 (0.7-2.7) | Triangular | 8.1 (2.3-13.8) | Triangular |
| **EFs of $CH_4$ (g $CO_2$-eq/$m^3$)** | **Distribution** | **EFs of $N_2O$ (g $CO_2$-eq/$m^3$)** | **Distribution** |
| 13.1 (7.7-28.1) | Triangular | 61.4 (23.3-184.9) | Triangular |

**Note:** EFs of GHG ($CH_4$ and $N_2O$) of wastewater treatment by $A^2O$ technology are extensively investigated from references[3-14], and values in brackets represent range within the 95% confidence intervals.

**Supplementary Table S11 | Market prices of electricity and chemicals consumed in wastewater treatment and nano-CdS.**

| Terms | Baseline | Unit | Distribution |
|---|---|---|---|
| Electricity | 0.070 | $/kWh | Triangular [a] |
| Acetic acid | 0.62 | $/kg | Triangular [c] |
| Polyacrylamide | 1.97 | $/kg | Triangular [c] |
| Lime | 0.26 | $/kg | Triangular [b] |
| Sand | 7.24E-03 | $/kg | Triangular [c] |
| Gravel | 5.57E-03 | $/kg | Triangular [c] |
| Aluminum chloride | 0.066 | $/kg | Triangular [c] |
| Hydrochloric acid | 0.14 | $/kg | Triangular [c] |
| Buffer solution | 0.06 | $/L | Triangular [c] |
| Nano-CdS | 99.26 | $/kg | Triangular [d] |

**Note: a, b, d,** distribution of market prices is provided in Table S13. **c,** distribution of market prices are assumed with fluctuation of ±25% upon the baseline value. Calculation of economic cost of conventional wastewater is according to an extended life-cycle economic evaluation (LCE) method based on LCA method, which has similar steps with those of LCA. Within the goal and scope defined by LCA, LCE was used to calculate the economic burdens in the whole life cycle process and identify the main links. Unit price of each item was taken from the average price of global market, and the average value were used. The cash flow and discount rate are not considered as a static analysis method of LCE as generally conducted in this research.

**Supplementary Table S12 | Estimated unit offset of GHG emission and economic cost from wastewater treatment and valuable by-products, nano-CdS.**

| Terms | Offset values | Unit | Distribution |
|---|---|---|---|
| Wastewater treatment | 0.95 | kg $CO_2$-eq/$m^3$ | Triangular |
| Wastewater treatment | 0.050 | \$/$m^3$ | Triangular |
| By-products (nano-CdS) | 99.26 | \$/kg | Triangular |

**Note:** Offset values provided here are baseline level, values of distribution including minimum, median and maximum following triangular distribution are summarized in the following Table S13.

**Supplementary Table S13 | Parameters and probability distributions used in the illustrative routes.**

| Parameters | Unit | Baseline | Distributions | References/Sources |
|---|---|---|---|---|
| GHG of partial oxidation | kg $CO_2$-eq/kg acetylene | 0.12 | 0.10, 0.12, 0.14, triangular | [15] |
| 1-butanol loss | kg/kg 2,3-butaediol | 0.028 | 0.025, 0.028, 0.030, triangular | [16,17] |
| Natural gas price | $/kg | 0.253 | 0.198, 0.253, 0.304, triangular | [18] |
| Electricity price | $/kWh | 0.070 | 0.067, 0.070, 0.074, triangular | Global market |
| Methanol price | $/kg | 0.50 | 0.48, 0.50, 0.56, triangular | Global market |
| Oxygen (liquid) price | $/kg | 0.058 | 0.057, 0.058, 0.059, triangular | Global market |
| N-methylpyrrolidone price | $/kg | 3.52 | 3.45, 3.52, 3.69, triangular | Global market |
| Hydrogen price | $/kg | 4.91 | 4.85, 4.91, 5.14, triangular | [16] |
| Industrial water price | $10^{-4}$ $/kg | 4.40 | 3.96, 4.40, 4.84, triangular | [19] |
| Sulfuric acid price | $/kg | 0.095 | 0.091, 0.095, 0.10, triangular | [20,21] |
| Caustic soda price | $/kg | 0.37 | 0.28, 0.37, 0.46, triangular | Global market |
| Quicklime price | $/kg | 0.26 | 0.16, 0.26, 0.29, triangular | [20,22] |
| Corn stover price | $/kg (dry) | 0.071 | 0.060, 0.071, 0.084, triangular | [20,23] |
| Cellulase price | $/kg | 2.88 | 2.59, 2.88, 3.45, triangular | [19] |
| DAP price | $/kg | 1.48 | 1.44, 1.48, 1.55, triangular | Global market |
| CSL price | $/kg | 0.093 | 0.086, 0.093, 0.100. triangular | Global market |
| Yeast price | $/kg | 1.44 | 1.12, 1.44, 1.80, triangular | Global market |
| 1-butanol price | $/kg | 1.15 | 1.10, 1.15, 1.24, triangular | Global market |
| 1,4-butanediol price | $/kg | 1.44 | 1.36, 1.44, 1.79, triangular | Global market |
| Nano-CdS price | $/kg | 99.26 | 97.82, 99.26, 100.70, triangular | Global market |
| COD of influent (WW) | $kg/m^3$ | 0.31 | 0.21, 0.30, 0.51, triangular | Estimation |
| TN of influent (WW) | $kg/m^3$ | 0.026 | 0.016, 0.026, 0.036, triangular | Estimation |
| TSS of influent (WW) | $kg/m^3$ | 0.11 | 0.050, 0.11, 0.17, triangular | Estimation |
| COD removal by WWT | % | 93 | 90, 93, 96, triangular | Estimation |
| TN removal by WWT | % | 70 | 60, 70, 80, triangular | Estimation |
| TSS removal by WWT | % | 94 | 92, 94, 96, triangular | Estimation |
| EFs of $CH_4$ (WWT) | $10^{-3}$ kg $CH_4$/kg COD | 1.7 | 0.7, 1.7, 2.7, triangular | [3-14] |
| EFs of $N_2O$ (WWT) | $10^{-3}$ kg $N_2O$/kg TN | 8.1 | 2.3, 8.1, 13.8, triangular | [3-14] |
| EFs of $CH_4$ (WWT) | g $CO_2$-eq/$m^3$ | 13.1 | 7.7, 13.1, 28.1, triangular | [3-14] |
| EFs of $N_2O$ (WWT) | g $CO_2$-eq/$m^3$ | 61.4 | 23.3, 61.4, 184.9, triangular | [3-14] |
| Acetic acid price | $/kg | 0.62 | 0.46, 0.62, 0.77, triangular | Global market |
| Polyacrylamide price | $/kg | 1.97 | 1.48, 1.97, 2.46, triangular | Global market |
| Sand price | $10^{-3}$ $/kg | 7.24 | 5.43, 7.24, 9.05, triangular | Global market |
| Gravel price | $10^{-3}$ $/kg | 5.57 | 4.18, 5.57, 6.96, triangular | Global market |
| Aluminum chloride price | $/kg | 0.066 | 0.049, 0.066, 0.082, triangular | Global market |
| Caustic soda price | $/kg | 0.25 | 0.20, 0.25, 0.30, triangular | Global market |
| Hydrochloric acid price | $/kg | 0.14 | 0.10, 0.14, 0.18, triangular | Global market |
| Buffer solution price | $/L | 0.06 | 0.04, 0.06, 0.08, triangular | Global market |
| Compressed air price | $/$m^3$ | 0.026 | 0.022, 0.026, 0.030, triangular | Global market |
| COD of influent (substrate) | $kg/m^3$ | 85 | 80, 85, 90, triangular | Lab estimation |
| TN of influent (substrate) | $kg/m^3$ | 4.5 | 4.0, 4.5, 5.0, triangular | Lab estimation |
| HM ions of influent (substrate) | $kg/m^3$ | 0.011 | 0.008, 0.011, 0.014, triangular | Lab estimation |
| Sugars content of molasses waste | % | 24 | 20, 24, 28, triangular | Industrial estimation |
| COD removal by SDBS | % | 70 | 28, 32, 36, triangular | Lab estimation |
| TN removal by SDBS | % | 15 | 10, 15, 20, triangular | Lab estimation |
| HM ions removal by SDBS | % | 96 | 94, 96, 98, triangular | Lab estimation |
| pH of substrate | —— | 7.00 | 6.95, 7.00, 7.05, triangular | Lab estimation |
| 2,3-BDO yield | g/L | 13 | 11, 13, 15, triangular | Lab estimation |
| Nano-CdS yield | (dry-weight) g | 0.04 | 0.02, 0.04, 0.06, triangular | Lab estimation |

**Note:** Parameters affecting the net GHG emission and economic burdens of 1,4-butanediol production by fossil-fuels refining, 2,3-butanediol production by bio-fermentation and solar-driven biohybrid synthesis, and conventional treatment of industrial wastewater are summarized here to comprehensively analyze the uncertainty of the aforementioned LCA models in this research. DAP, diammonium phosphate; CSL, Corn steep liquor; CdS, cadmium sulfide; COD, chemical oxygen demand; TN, total nitrogen; TSS, total suspended solids; EFs, emission factors; WW, wastewater; WWT, wastewater treatment; HM, heavy metals; SDBS, solar-driven biohybrid synthesis; 2,3-BDO, 2,3-butanediol.

### Supplementary Figures

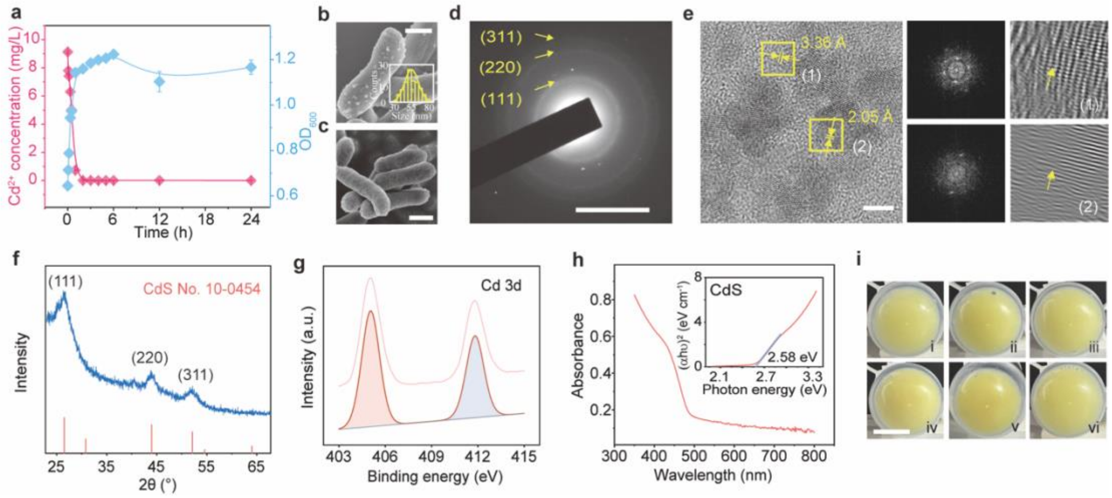


**Supplementary Figure S1** | Other evidences for the production of CdS semiconductor biohybrid. **a**) Time courses of $Cd^{2+}$ removal and $OD_{600}$ of *V. natriegens* XG203 cultured in a well-defined minimum medium with addition of $Cd^{2+}$ (0.1 mM). Error bar: standard deviation, n≥3. **b**) SEM image of semiconductor biohybrid collecting from **a**) at 24 h. SEM images are used to calculate the size distribution of nanoparticles (more than 110 particles). Scale bar, 0.5 µm. **c**) SEM image of strain XG203 only cultured in a well-defined minimum medium. Scale bar, 0.5 µm. **d**) SAED pattern of CdS nanoparticles. Nanoparticles were lysed from biohybrid. Scale bar, 5 1/nm. **e**) HRTEM and FFT images of CdS nanoparticles. Scale bar, 5 nm. **f**) XRD patterns of CdS nanoparticles. XRD patterns are in line with that of crystalline cubic CdS (JCPDS No. 10-0454). Three main characteristics peaks are attributed to crystal plane (111), (220), and (311), corresponding to lattice fringes with the spacing of 3.36, 2.05, 1.75 Å, respectively. **g**) High-resolution XPS spectra of Cd 3d in the nanoparticles of biohybrid. **h**) UV-Vis absorption spectra of CdS nanoparticles. Direct band energy is calculated in Tauc plots (inset) according to UV-Vis spectra. **i**) Visual representation of in-situ production of semiconductor biohybrids using common wastewater organics. Strain XG203 was cultured in a well-defined minimum medium with addition of $Cd^{2+}$ (0.1 mM) and carbon sources from common wastewater organics, including i) glucose, ii) fructose, iii) sucrose, iv) maltose, v) starch, vi) acetate, respectively. Scale bar, 1.5 cm.

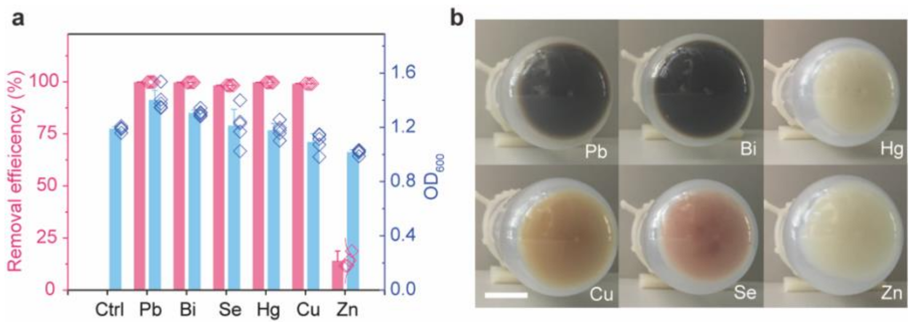


**Supplementary Figure S2 | Bacterial capability for the production of various semiconductor biohybrids using different heavy metals.** **a**) The removal of heavy metal by *V. natriegens* XG203 cultured in a well-defined minimum medium with addition of one kind of heavy metal. The concentration of all metal ions were 0.1 mM, but the $Hg^{2+}$ was 0.01 mM. The removal efficiency is higher than 98% for Pb, Bi, Cu, Se, and Hg, and 14.04% for Zn. Error bar: standard deviation, n≥4. **b**) Visual representation of pellets collecting from **a**). Colors potentially indicate the production of semiconductor biohybrids. Scale bar, 1.2 cm.

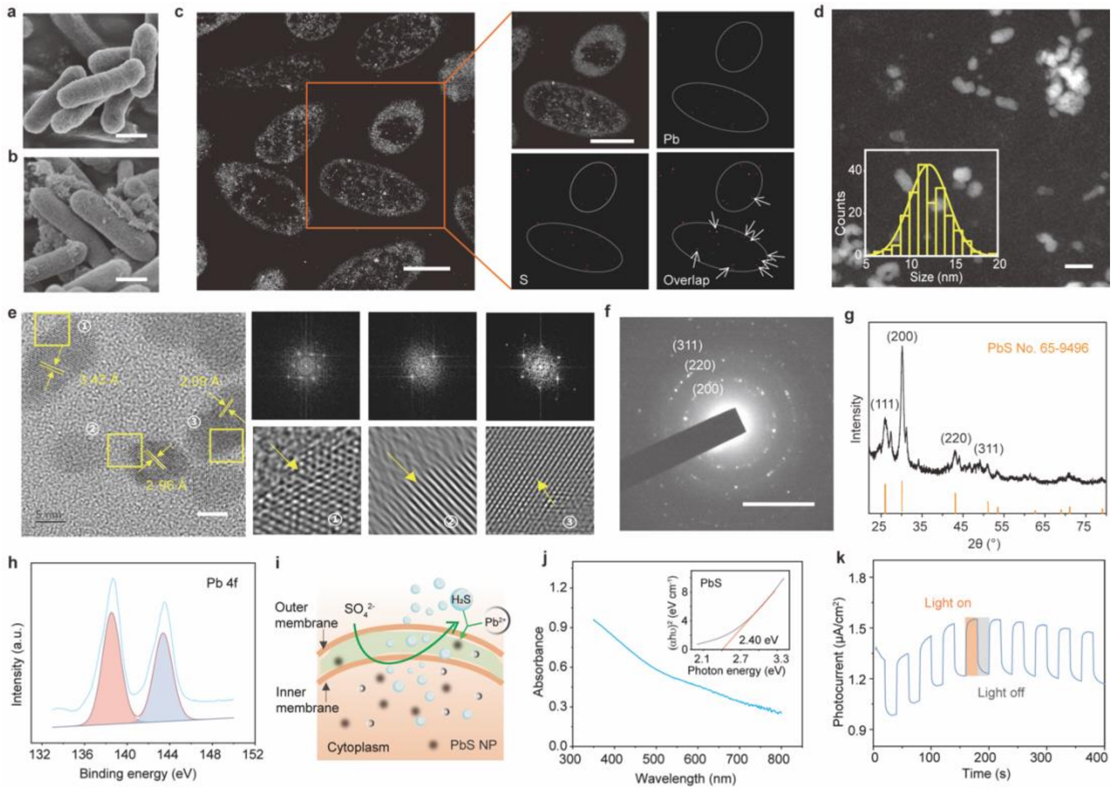


**Supplementary Figure S3 |** Production of PbS semiconductor biohybrid. **a**) SEM image of *V. natriegens* XG203 cultured in a well-defined minimum medium. Scale bar, 0.5 μm. **b**) SEM image of biohybrid samples collecting from pellets of strain XG203 cultured in a well-defined minimum medium with addition of $Pb^{2+}$ (0.1 mM) in **Fig. S2a**). Scale bar, 0.5 μm. **c**) Cross-sectional STEM and EDS mapping images of biohybrid samples. Nanoparticles are distributed inside the bacterial cell and composed of Pb and S elements. Scale bars, 0.5 μm. **d**) HAADF-STEM images of PbS nanoparticles. Nanoparticles were lysed from biohybrid samples. PbS nanoparticles have an average size of ~12.20 nm. HRTEM images are used to calculate the size distribution of PbS nanoparticles (more than 200 particles). Scale bar, 20 nm. **e**) HRTEM and FFT images of PbS nanoparticles. PbS nanoparticles show the lattice spacing of 3.42, 2.96, 2.09 Å and the local crystallization. Scale bar, 5 nm. **f**) SAED patterns of PbS nanoparticles. PbS nanoparticles show the crystal pane (200), (220), and (311). Scale bar, 5 1/nm. **g**) XRD patterns of PbS nanoparticles. XRD patterns are in line with that of crystalline PbS (JCPDS No. 65-9496). Four main characteristics peaks correspond to crystal plane (111), (200), (220), and (311), corresponding to lattice fringes with the spacing of 3.42, 2.96, 2.09, 1.79 Å, respectively. **h**) High-resolution XPS spectra of Pb 4f in biohybrid samples. Pb(II) is the main Pb species in the nanoparticles. **i**) Schematic diagram of *in-situ* production of semiconductor biohybrid. **j**) UV-Vis absorption spectra of PbS nanoparticles. Direct band energy of PbS nanoparticles is about 2.40 eV in Tauc plots (inset) calculated according to UV-Vis spectra. **k**) A representative photocurrent curve of isolated nanoparticles with light on/off cycles (20 s on and off, 100 mW/cm$^2$). The isolated nanoparticles were lysed from biohybrid samples.

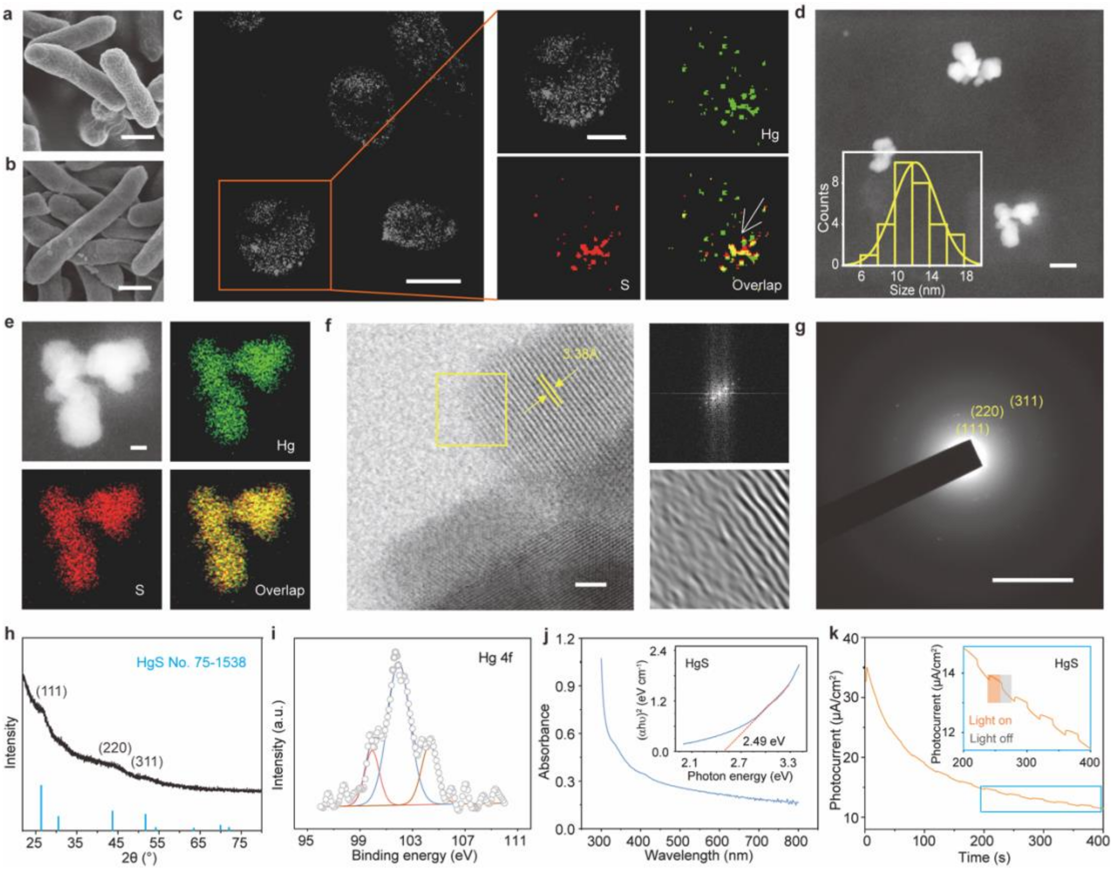


**Supplementary Figure S4** | Production of HgS semiconductor biohybrid. **a**) SEM image of strain XG203 cultured in a well-defined minimum medium. Scale bar, 0.5 μm. **b**) SEM image of biohybrid samples collecting from pellets of strain XG203 cultured in a well-defined minimum medium with addition of $Hg^{2+}$ (0.01 mM) in **Fig. S2a**). Scale bar, 0.5 μm. **c**) Cross-sectional STEM and EDS mapping images of biohybrid samples. Nanoparticles are mainly distributed inside the bacterial cell and composed of Hg and S elements. Scale bars, 0.5 μm (left) and 0.3 μm (right). **d**) HAADF-STEM images of HgS nanoparticles. HgS nanoparticles were lysed from biohybrid samples. Nanoparticles have an average size of ~12.38 nm. HRTEM images are used to calculate the size distribution of nanoparticles (more than 30 particles). Scale bar, 20 nm. **e**) HAADF-STEM and EDS mapping images of isolated HgS nanoparticles. HgS nanoparticles are composed of Hg and S elements. Scale bar, 5 nm. **f**) HRTEM and FFT images of HgS nanoparticles. HgS nanoparticles show the main crystal plane (111). Scale bar, 2 nm. **g**) SAED patterns of isolated HgS nanoparticles. Nanoparticles show the crystal pane (111), (220), and (311). Scale bar, 5 1/nm. **h**) XRD patterns of HgS nanoparticles. XRD patterns are in line with that of crystalline cubic HgS (JCPDS No. 75-1538). Three main characteristics peaks are attributed to crystal plane (111), (220), and (311), corresponding to lattice fringes with the spacing of 3.38, 2.07, 1.77 Å, respectively. **i**) High-resolution XPS spectra of Hg 4f in biohybrid samples. Hg(II) is the Hg species in the nanoparticles. **j**) UV-Vis absorption spectra of HgS nanoparticles. Direct band energy of HgS nanoparticles is about 2.49 eV in Tauc plots (inset) calculated according to UV-Vis spectra. **k**) A representative photocurrent curve of isolated nanoparticles with light on/off cycles (20 s on and off, 100 mW/$cm^2$). The isolated nanoparticles were lysed from biohybrid samples.

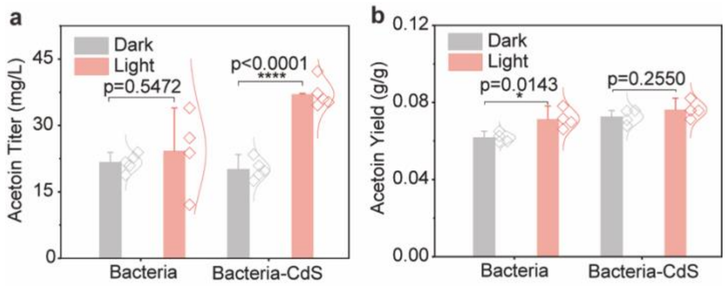


**Supplementary Figure S5 | ** Solar-driven production of acetoin, a direct precursor for BDO production. **a**) Production of acetoin by the prepared semiconductor biohybrid using a well-defined minimum medium with addition of glucose under dark and light condition (4.2 $mW/cm^2$). Error bar: standard deviation, n≥4. **b**) Waste sugar-to-acetoin conversion yield by the *in-situ* production of semiconductor biohybrid using wastewater medium under dark and light condition (4.2 $mW/cm^2$). Error bar: standard deviation, n=4. P values are determined by a two-tailed unpaired t-test.

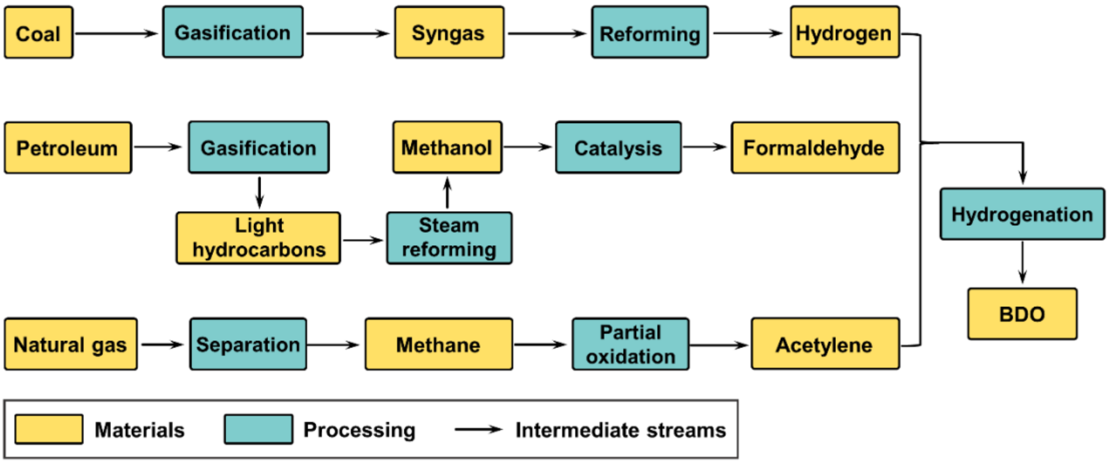


**Supplementary Figure S6** | Schematic and system boundary of 1,4-butanediol production by fossil-fuels refining. Feedstocks of formaldehyde (derives from methanol production by the steam reforming of light hydrocarbons and syngas) and gaseous acetylene (derives from partial oxidation of methane) are catalyzed into intermediate butyne-diol via Reppe process, which is hydrogenated to 1,4-butanediol.

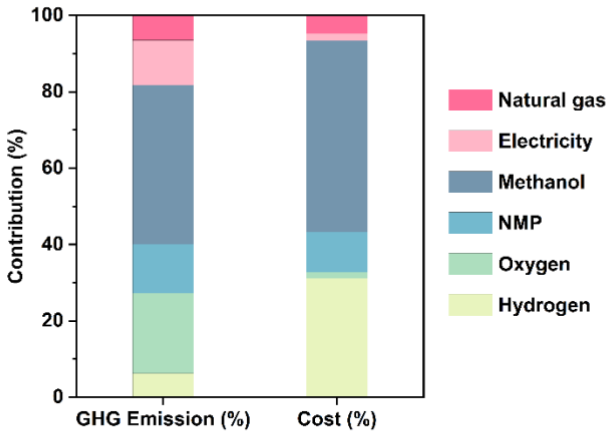


**Supplementary Figure S7** | Contributions of factors to GHG emission and cost of 1,4-butanediol production by fossil-fuels refining. In the synthesis processes, feedstocks of methanol contribute to the largest GHG emission (41.63%) and cost (50.23%) of fossil-fuel refining.

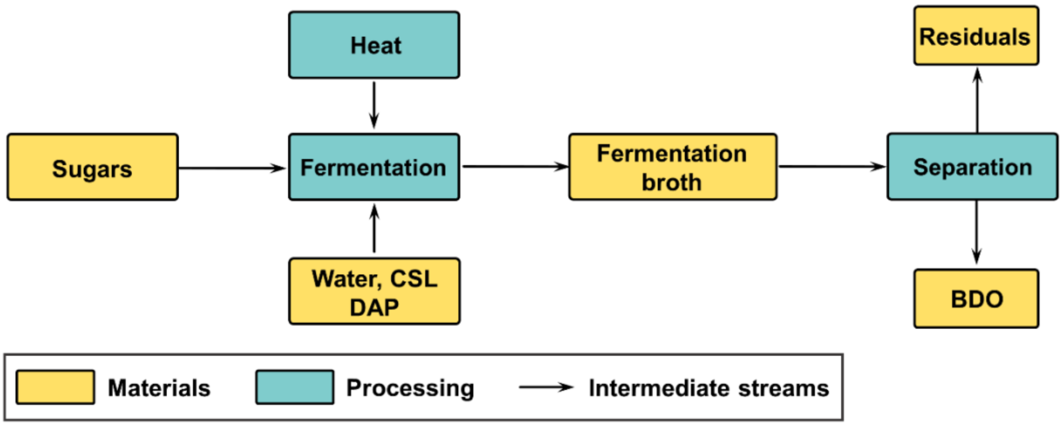


**Supplementary Figure S8 |** Schematic and system boundary of 2,3-butanediol production by fermentation of sugars. Slurry of sugars is pre-heated (121 °C, ~15 min) and fermented at 37 °C with 48 hours residence. 2,3-butanediol is separated from the fermentation broth via sequential flash-gas purging, centrifugation, and hybrid extraction-distillation.

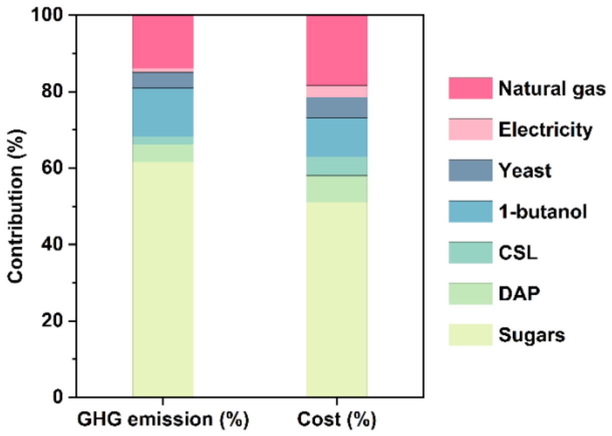


**Supplementary Figure S9** | Contributions of factors to GHG emission and cost of 1,4-butanediol production by fermentation of sugars. Sugars dominate the GHG emission (61.66%) and cost (51.03%) of sugars fermentation.

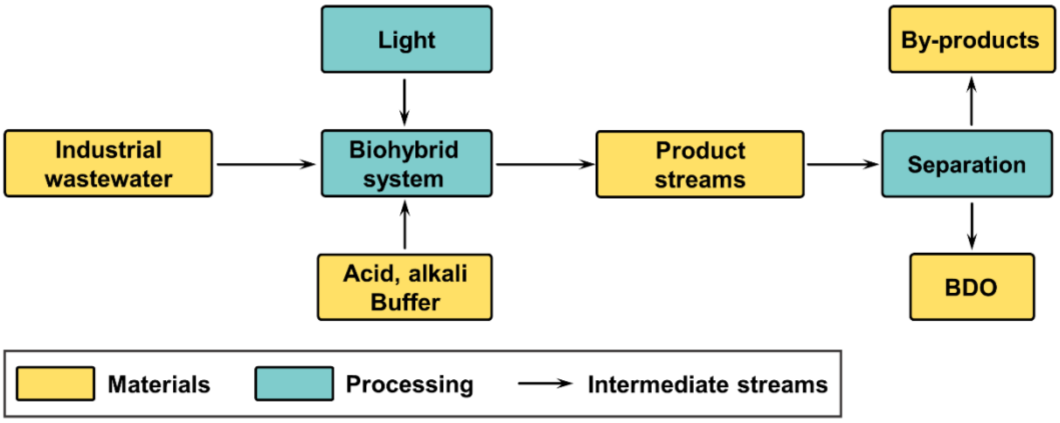


**Supplementary Figure S10 |** Schematic and system boundary of 2,3-butanediol production by solar-driven biohybrid using wastewater. Lab-scale production of 2,3-butanediol by solar-driven biohybrid using wastewater is assumed to be modified with an industrial modelling similar to bio-fermentation, including culture of *V. natriegens* using wastewater medium in an illuminated fermenter, pH adjustment by hydrochloric acid and caustic soda with buffer solution, and separation of products and by-products.

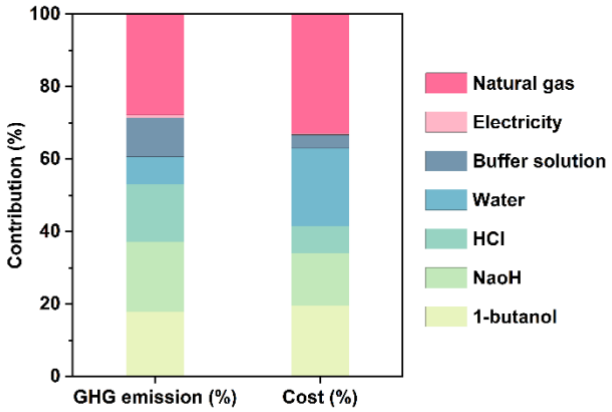


**Supplementary Figure S11** | Contributions of factors to GHG emission and cost of 2,3-butanediol production by solar-driven biohybrid using wastewater. The energy (natural gas) consumption for products separation from culture systems accounts for 28.66% of GHG emission and 33.31% of cost. The costly feedstocks were replaced by the cost-free wastewater to decouple 2,3-butanediol production from commodity prices.

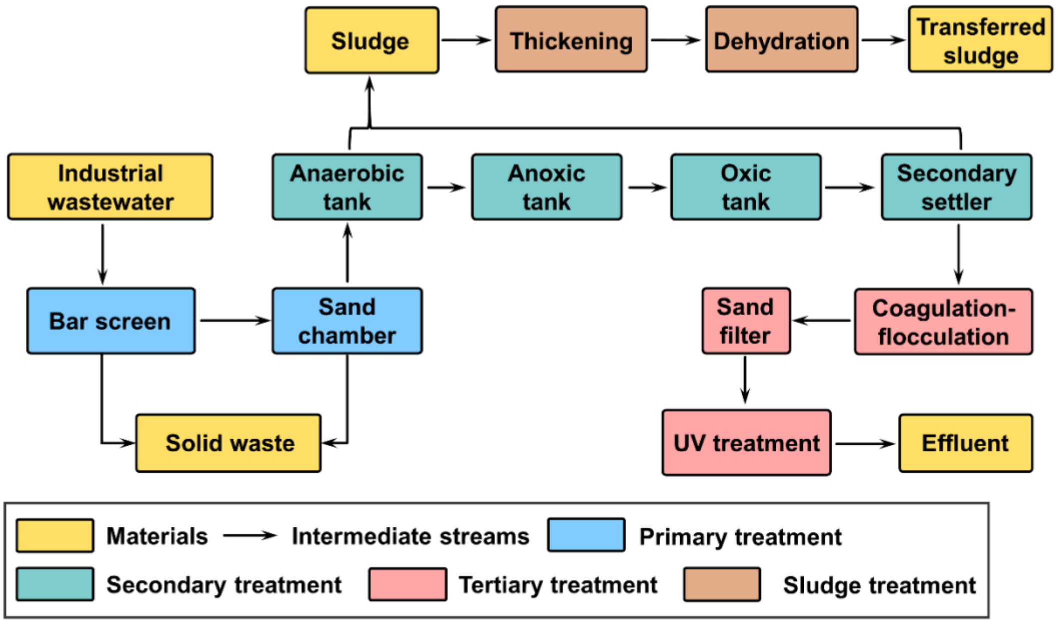


**Supplementary Figure S12** | Schematic and system boundary of conventional wastewater treatment process used for treating industrial wastewater with high COD and heavy metals. In the typical wastewater treatment process, anaerobic-anoxic-aerobic ($A^2O$) as the secondary treatment with front-end primary treatment is assumed here to remove high-strength COD in the molasses wastewater. The coagulation/flocculation-sedimentation-filtration-ultraviolet reduction as the tertiary treatment is assumed here to concentrate heavy metal ions in the electroplating wastewater. The residual sludge of wastewater treatment is thickened, dewatered, and transferred to the landfill sites.
